# KIF1C activates and extends dynein movement through the FHF cargo adaptor

**DOI:** 10.1101/2023.10.26.564242

**Authors:** Ferdos Abid Ali, Alexander J. Zwetsloot, Caroline E. Stone, Tomos E. Morgan, Richard F. Wademan, Andrew P. Carter, Anne Straube

## Abstract

Cellular cargos move bidirectionally on microtubules due to the presence of opposite polarity motors dynein and kinesin. Many studies show these motors are co-dependent, whereby one requires the activity of the other, although the mechanism is unknown. Here, using in vitro motility assays, we show that the kinesin-3 KIF1C acts both as an activator and a processivity factor for dynein. Activation only requires a fragment of the non-motor tail of KIF1C (KIF1C-stalk) to bind the cargo adaptor HOOK3. Cryo-EM, crosslinking mass spectrometry and AlphaFold2 predictions reveal this binding site to be separate from that of two constitutive factors (FTS and FHIP), which link HOOK3 to small G-proteins on cargos. We provide a structural model for how the FTS-HOOK3-FHIP1B (FHF) complex is auto-inhibited and explain how the KIF1C-stalk relieves this inhibition. Collectively, our work provides a molecular explanation for co-dependency by revealing that the mutual activation of dynein and kinesin is mediated through their shared adaptor. Many adaptors bind both dynein and kinesins, suggesting this mechanism could be generalised to other bidirectional complexes.

## INTRODUCTION

The organisation of a cell’s content is central to its function. Long-range transport of membrane organelles depends on the opposing microtubule-based motors dynein and kinesin. Almost all cellular cargos move back and forth due to the simultaneous presence of both types of motor, a concept known as bidirectional transport^1–3^. Such an arrangement requires coordination between kinesin and dynein to prevent unproductive stalling and tug-of-war. A large body of evidence shows that opposite polarity motors also depend on each other during motility; diminishing either dynein or kinesin-1 activity leads to predominantly static organelles, lipid droplets and mRNA-protein complexes^4–11^. This phenomenon, referred to as co-dependence of antagonistic motors^2^, also occurs between dynein and kinesin-3s. For example, depletion of KIF1C reduces transport of integrin-containing vesicles to the same extent in both directions^12^ and the retrograde transport of dense core vesicles is perturbed when the *C. elegans* KIF1A homologue Unc-104 is mutated^12,13^. Although it is well established that dynein and kinesin require each other on cargos, the molecular mechanism of co-dependence remains unclear.

## RESULTS

### Dynein and KIF1C form a co-dependent complex scaffolded by HOOK3

Human cytoplasmic dynein is a weakly processive motor on its own^14^, requiring a co-factor dynactin and a coiled-coil cargo adaptor for full activation^15– 17^. Many of dynein’s activating cargo adaptors can also bind kinesins, including BICD2^18,19^, BICDR1^20,21^, HOOK^21–25^, TRAK^26,27^ and JIP3^28–30^. HOOK3 is of particular interest as it has been recently shown to simultaneously recruit dynein/dynactin and the kinesin-3 motor KIF1C *in vitro*^22^. To determine whether these opposite polarity motors show co-dependence, we reconstituted purified dynein/dynactin-HOOK3-KIF1C (DDHK) and analysed their motile behaviour using single molecule total internal reflection fluorescence (TIRF) microscope assays.

In the presence of all components, complexes undertook processive minus end (dynein-driven, 80%) or plus end-directed transport (kinesin-driven, 16%) (Fig. 1a-b). The observation of predominant minus end runs agrees with a previous study on this co-complex^22^. In addition, we observe occasional switches in direction during runs (2%) (Fig. 1b-c), reminiscent of bidirectional cargo movement in the cell^24^. This suggests that coupling opposite polarity motors via their adaptors supports processive unidirectional transport and prevents tug-of-war. This differs from artificial, DNA-linked systems where the motors resist each other’s movement^31–33^.

**Figure 1:**
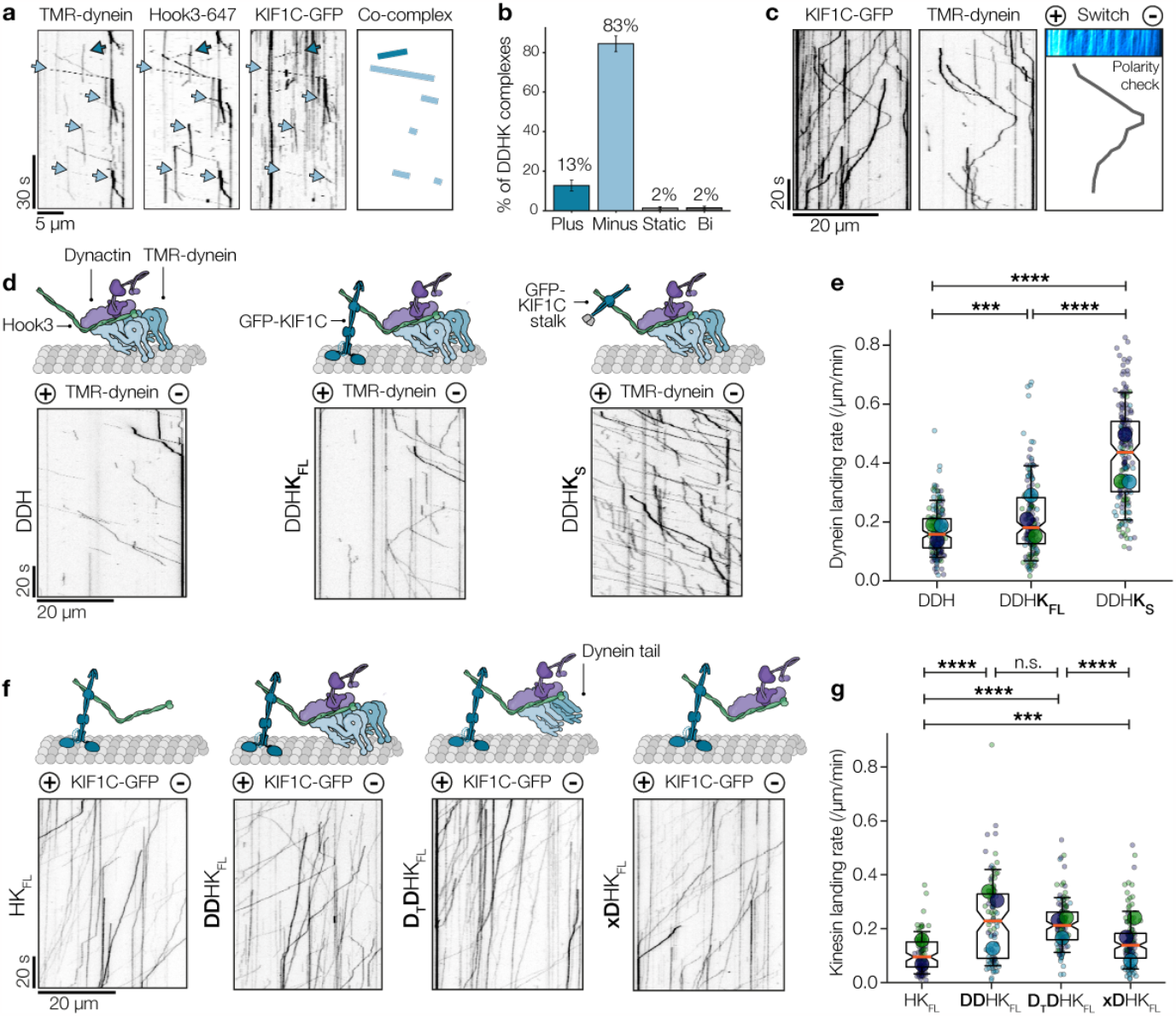
Dynein and KIF1C are co-dependent in vitro. **(a)** Kymographs show co-localised movement of TMR-dynein, HOOK3-647 and KIF1C-GFP along microtubules. Right panel shows minus-end runs of DDHK complexes in light blue and plus-end runs in dark blue. **(b)** Quantification of DDHK complex directionality. n=1418 motor complexes. **(c)** Example of a directional switch of DDHK complex during a run. Inset kymographs shows polarity check with KIF1C-GFP only, which was performed after every experiment containing both KIF1C and Dynein to establish microtubule polarity. **(d)** Schematics and representative kymographs of TMR-labelled dynein mixed with dynactin and HOOK3 (DDH) or dynactin, HOOK3 and full-length KIF1C-GFP (DDHK) or dynactin, HOOK3 and GST-Kif1Cstalk-GFP (DDHKS). All experiments performed in the presence of Lis1. **(e)** Superplot of TMR-dynein landing rate on microtubules for different co-complex combinations (as in d). Small dots indicate single microtubules and large dots indicate experimental averages. n=150-171 microtubules **(f)** Schematics and representative kymographs of KIF1C-GFP mixed with HOOK3 (HKFL), dynactin, HOOK3 and full-length dynein (DDHK), dynein tail, dynactin, HOOK3 and KIF1C-GFP (DTDHKFL) or dynactin, HOOK3 and KIF1C-GFP (xDHKFL). **(g)** Superplot of KIF1C-GFP landing rate on microtubules for different co-complex combinations (as in f). Small dots indicate single microtubules and large dots indicate experimental averages. n=94-129 microtubules. n.s. p>0.05, ***p<0.001, ****p<0.0001 Kruskal Wallis H test followed by a Conover’s posthoc test to evaluate pairwise interactions with a multiple comparison correction applied using Holm–Bonferroni.

We noticed a significant increase in the dynein landing rate in the presence of full-length KIF1C (KFL) (Fig. 1d-e), an effect which has not been previously described. This is mediated through HOOK3 and dynactin as omitting either of these factors abolishes dynein motility (Extended Data Fig. 1). To dissect the mechanism of this activation, we focused on a KIF1C stalk construct (GST-KIF1C^642-922^-GFP, KS) that does not contain the motor domain but includes the region required to bind HOOK3^23^. Strikingly, dynein microtubule recruitment increases 2.6-fold in the presence of the KIF1C stalk construct (Fig. 1d-e), indicating that this region is sufficient for binding HOOK3 and activating the co-complex. A similar activating effect was observed vice versa; KIF1C landing rate on microtubules increased two-fold in the presence of dynein and dynactin (DDHK) compared to HOOK3-KIF1C (HKFL), even if the motor domain was omitted by using a dynein tail construct (DT) (Fig. 1f-g). Dynactin alone is also sufficient to increase the KIF1C-HOOK3 landing rate, although to a lesser extent (1.4-fold compared to KIF1C-HOOK3) (Fig. 1f-g). Collectively, our results show that KIF1C and dynein can facilitate each other’s microtubule binding activity using their non-motor domains.

### FHF is an auto-inhibited cargo adaptor

In order to understand how this mutual activation is mediated, we investigated the regulation of the cargo adaptor. Unlike other cargo adaptors, HOOKs form a stable assembly with two other accessory factors, FTS and FHIP (FHF complex) (Extended Data Fig. 2a)^34,35^. However, the proposed binding sites of KIF1C and FTS-FHIP1B to HOOK3 are currently unclear beyond the adaptor’s C-terminal 165 residues^22^. To address its unknown molecular organisation, we determined the structure of FHF by cryo-EM. Our 3.2 Å reconstruction of full-length FTS and FHIP1B in complex with a C-terminal HOOK3 fragment (residues 571-718) allowed an atomic model to be built for each subunit (Fig. 2a-b, Extended Data Fig. 2b, Extended Data Fig. 3). The cryo-EM structure shows that HOOK3 is integral to the complex (Fig. 2b), explaining why FTS and FHIP cannot form a subcomplex on their own over gel filtration (Extended Data Fig. 2c). The interactions involve the last of HOOK3’s four coiled-coil domains (CC4) and a C-terminal extension (CTE), consisting of a flexible region followed by a single alpha helix (residues 694-702) (Fig. 2b, Extended Data Fig. 3g). The first anchor point (site 1) involves one of the CTEs, with the helix latching onto a hydrophobic pocket within FHIP1B. Site 2 comprises a span of HOOK3 CC4 residues (653-688), packing tightly against the FTS catalytically dead ubiquitin ligase domain (Fig. 2b, Extended Data Fig. 3h). At site 3, the opposite side of HOOK3 CC4 (spanning residues 663-670) binds a C-terminal alpha-helix on FTS. This helix is in a long loop (FTS “belt”) that wraps around HOOK3 before docking into a pocket on FHIP1B (FHIP “latch”) (Fig. 2b, Extended Data Fig. 3h-i). Together, these interactions mean that the extreme C-terminus of HOOK3 is buried within FHIP1B and FTS and cannot interact with cargo directly. Instead, the opposing face of FHIP1B comprises a highly conserved patch of surface-exposed residues (Extended Data Fig. 4a) that AlphaFold2 suggests is a binding site for small G-protein Rab5, which has been shown to link FHF to early endosomes (Fig. 2c and Extended Data Fig. 4b)^34,36^. The location of this site means the adaptor is positioned to be extended away from the cargo.

**Figure 2:**
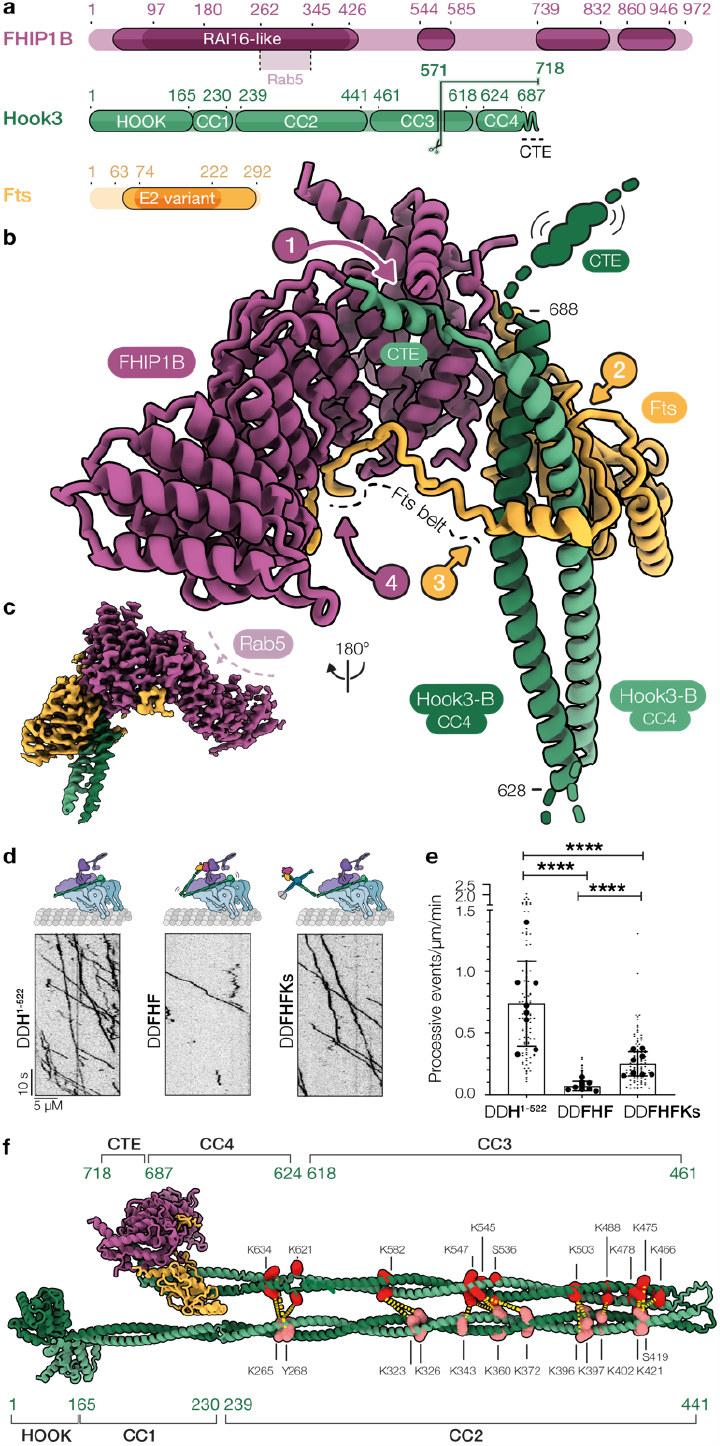
FHF is an auto-inhibited adaptor. **(a)** Domain architecture of FHIP1B, HOOK3 and FTS. Putative Rab5 binding site indicated in FHIP1B. Scissor cut at residue 571 in HOOK3 indicates truncation site used for FHF cryo-EM construct. (b) Atomic model of FHF complex built from 3.2 Å cryo-EM structure, highlighting the four main interaction sites and the flexible regions within HOOK3. **(c)** Cryo-EM density of FHF with AlphaFold2-predicted Rab5 binding site highlighted on FHIP1B. See Extended Data Fig. 4 for details. **(d)** Representative kymographs of TMR-dynein and dynactin in presence of either HOOK31-522 (left panel), full-length FHF (middle panel) or full-length FHF + KIF1C stalk (right panel). All samples also included Lis1. **(e)** Quantification of processive events per μm per min from 8 technical replicates. Bars indicate mean ± S.D., small dots indicate data for each microtubule, large dots indicate experimental averages. n=120 microtubules (15 per replicate). ****=p<0.0001 (Kruskal-Wallis non-parametric test with Dunn’s multiple comparison). (f) Disuccinimidyl sulfoxide (DSSO) crosslinks (yellow dashes) between HOOK3N residues (salmon orange) and HOOK3C residues (red) mapped onto a stitched AlphaFold2 model of full-length FHF (see methods).

Factors that bind the C-terminus of other adaptors, such as BICD2 and Spindly, are thought to activate them^37–41^. To investigate the activity of the full-length FHF cargo adaptor, we tested its ability to facilitate processive dynein movement in single molecule motility assays. We found that processive dynein-dynactin-FHF (DDFHF) complexes were formed 11-fold less efficiently than DDH complexes made with a constitutively active truncated HOOK3 construct (DDH^1-522^) (0.065 vs 0.75 processive events/µm/s, respectively) (Fig. 2d-e). These data suggest that FHF is in an auto-inhibited conformation and that FTS-FHIP1B binding do not activate HOOK3. In contrast, adding KIF1C stalk led to a ~3.7-fold increase in processive events compared to a DDFHF control (Fig. 2d-e). This confirms that KIF1C stalk activates HOOK3 also in the context of FHF.

To understand the FHF auto-inhibition at the structural level, we used cross-linking mass spectrometry and mapped the interactions within the full-length molecule. We found numerous crosslinks between the HOOK3 coiled-coil domains, with CC2 being in close proximity to CC3 and CC4 (Extended Data Fig 5a). We mapped these crosslinks onto a model of full-length FHF, constructed by combining our cryo-EM model with three different and partially overlapping AlphaFold2 predicted segments of HOOK3 (Fig. 2f, Extended Data Fig 5b). The crosslinking is consistent with a folded-back conformation of HOOK3, in line with other adaptors such as BICD2 and Spindly^37–40^. This inhibited configuration clashes with dynactin/dynein tail binding to HOOK3, which is a pre-requisite for motor activation (Extended Data Fig 5b)^15–17,21^.

### KIF1C stalk binds FHF at a key regulatory site on HOOK3 to open up the adaptor

Next, we addressed how KIF1C stalk binds HOOK3 and interplays with FTS and FHIP1B at the molecular level. KIF1C is a stable dimer consisting of an N-terminal motor domain and a C-terminal tail. To obtain a full-length predicted structure of KIF1C, we stitched seven AlphaFold2 models together (Extended Data Fig 6a). Our structure suggests there are three coiled-coils (here called CC1-CC3) interspersed with two globular domains. The kinesin-3 specific forkhead-associated (FHA) domain sits between CC1 and CC2. The second region, between CC2 and CC3 was not previously known to be globular and we therefore refer to this whole folded region as the stalk globular domain (SGD) (Fig. 3a-b). The KIFC stalk we used in our experiments encompasses CC2, the SGD and CC3. Adding this KIF1C stalk to FHF in gel-filtration assays revealed that a stable complex can be reconstituted, indicating the binding sites for FTS-FHIP1B and KIF1C stalk do not overlap (Extended Data Fig. 7a). We used mass photometry to measure the molecular weight of the complex as ~434 kDa, which is close to the predicted molecular weight of one FHF and one dimeric KIF1C stalk (430 kDa) (Extended Data Fig. 7b).

**Figure 3.**
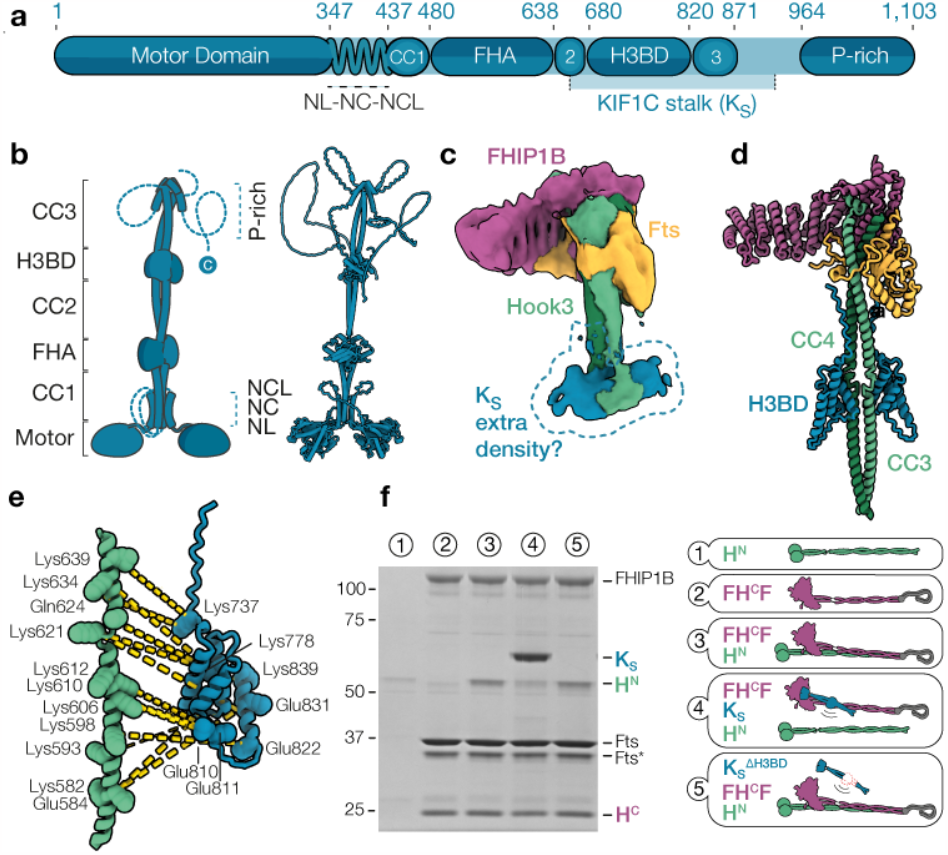
KIF1C stalk binds and activates FHF. (a) Domain architecture of KIF1C with the KIF1C stalk region highlighted below the main bar. FHA, Forhead-associated domain. SGD, Stalk globular domain. P-Rich, Proline-rich region. (b) Cartoon and stitched structural model of the AlphaFold2 predicted full-length KIF1C molecule. **c**, Segmented cryo-EM density of FHF bound to part of KIF1CS2 (674-922, blue dashed region), shown at low-threshold contour level. **d**, AlphaFold2 model of HOOK3 residues 571-718 bound to KIF1CSH3BD (722-840). H3BD, Hook3 binding domain. Note that that FTS, FHIP1B and HOOK3 (627-705) models from the cryo-EM structure are superposed on the HOOK3571-718-KIF1CSH3BD prediction to give a composite structure. **e**, A zoomed in inset from (d) showing one copy of the HOOK3 dimer at the CC3-CC4 junction bound to one copy of KIF1C H3BD dimer. Yellow dashes indicate crosslinks observed between the two proteins using DSSO crosslinker. (f) Pull-down assay using Strep-FHCF (comprising HOOK3 C-terminus 451-718), HN (HOOK3 N-terminus 1-450), Ks (GST-Kif1C stalk 642-922) and Ks ∆H3BD (GST-Kif1C stalk 642-922 ∆722-840). Key for the sample number above the gel is given in the right panel; colour-coded schematics indicate the presumed complexes formed. The grey fish hook schematic in FHCF cartoons represents the Strep-tag.

To map the binding sites more precisely, we used cryo-EM on a shorter KIF1C-stalk construct (674-922) bound to an FHF containing HOOK3 residues 451-718 (Extended Data Fig. 7c). We observed extra density corresponding to the KIF1C stalk at the hinge between HOOK3 CC3 and CC4 (Fig. 3c), although the resolution was low due to flexibility of this part of HOOK3 (Extended Data Fig. 6b, Extended Data Fig. 8a-c). We therefore used AlphaFold2 to screen pairwise interactions of sequential dimeric KIF1C fragments (in 120-residue increments) against a dimer of HOOK3 encompassing CC3 and CC4 (Extended Data Fig. 8d). This identified part of KIF1C’s SGD and CC3 as the top hit (residues 722-840, hereafter HOOK3 binding domain [H3BD]). The resulting AlphaFold2 model showed it binding HOOK3 at a position which matches the extra density in our cryo-EM structure (Fig. 3c, d). In further support of this assignment, deleting the H3BD from the KIF1C stalk abolished its interaction with HOOK3 (Extended Data Fig. 7d). The inverse is also true as FHF with a truncated HOOK3 (629-718), which lacks the kinesin binding site, no longer bound to the KIF1C stalk H3BD (Extended Data Fig. 7e). Furthermore, cross-linking mass spectrometry revealed numerous crosslinks from H3BD to the HOOK3 CC3-CC4 interface (Fig. 3e, Extended Data Fig. 8e). Overall our mapping studies showed that the KIF1C stalk H3BD mainly binds FHF within HOOK3 (residues 607-632), which is non-overlapping with the FTS and FHIP1B binding sites HOOK3 (residues 663-718).

We noted that KIF1C binding to HOOK3 is in close proximity to the inhibitory interaction between HOOK3’s CC2 and CC3-CC4 (Extended Data Fig. 8f). Therefore, KIF1C binding might disrupt the auto-inhibited form of FHF (Fig. 2f). To test this directly, we used pull-down experiments using a Streptagged FHF containing a C-terminal HOOK3 fragment (FH^C^F). In the absence of KIF1C stalk, this bound an N-terminal fragment of HOOK3 (H^N^) (Fig. 3f), which further supports our model that FHF is inhibited by CC2 folding back to contact CC3-CC4. When we added KIF1C stalk, we found it reduced H^N^ binding relative to background binding of H^N^ control. In contrast, a KIF1C stalk lacking the H3BD was not able to bind FHF or displace H^N^ (Fig. 3f). These data suggest that KIF1C stalk can shift the equilibrium towards an open configuration of HOOK3 and relieve FHF auto-inhibition.

### KIF1C acts as a processivity factor for dynein

Finally, we asked why it might be advantageous for KIF1C to bind dynein-dynactin-FHF. It could be a passive passenger using dynein to efficiently relocate to the cell centre, or alternatively could directly affect some aspect of dynein’s movement. To address this, we compared the motility parameters of complexes containing dynein, dynactin and HOOK3 to those co-assembled with either KIF1C stalk or full-length KIF1C (Fig. 4a). We found that the speed of minus-end directed runs is slightly decreased when the motor domain of KIF1C is present (Fig. 4b). This suggests that KIF1C engages with the microtubule when dynein drives the motility of the complex. We found both the duration motor complexes stay on the microtubule (“dwell time”) and the run length of dynein-driven motility increase by about 50% when the KIF1C motor is present (Fig. 4c-d). This suggests that KIF1C acts as a processivity factor for dynein and thereby stimulates the function of the opposite polarity motor.

**Figure 4:**
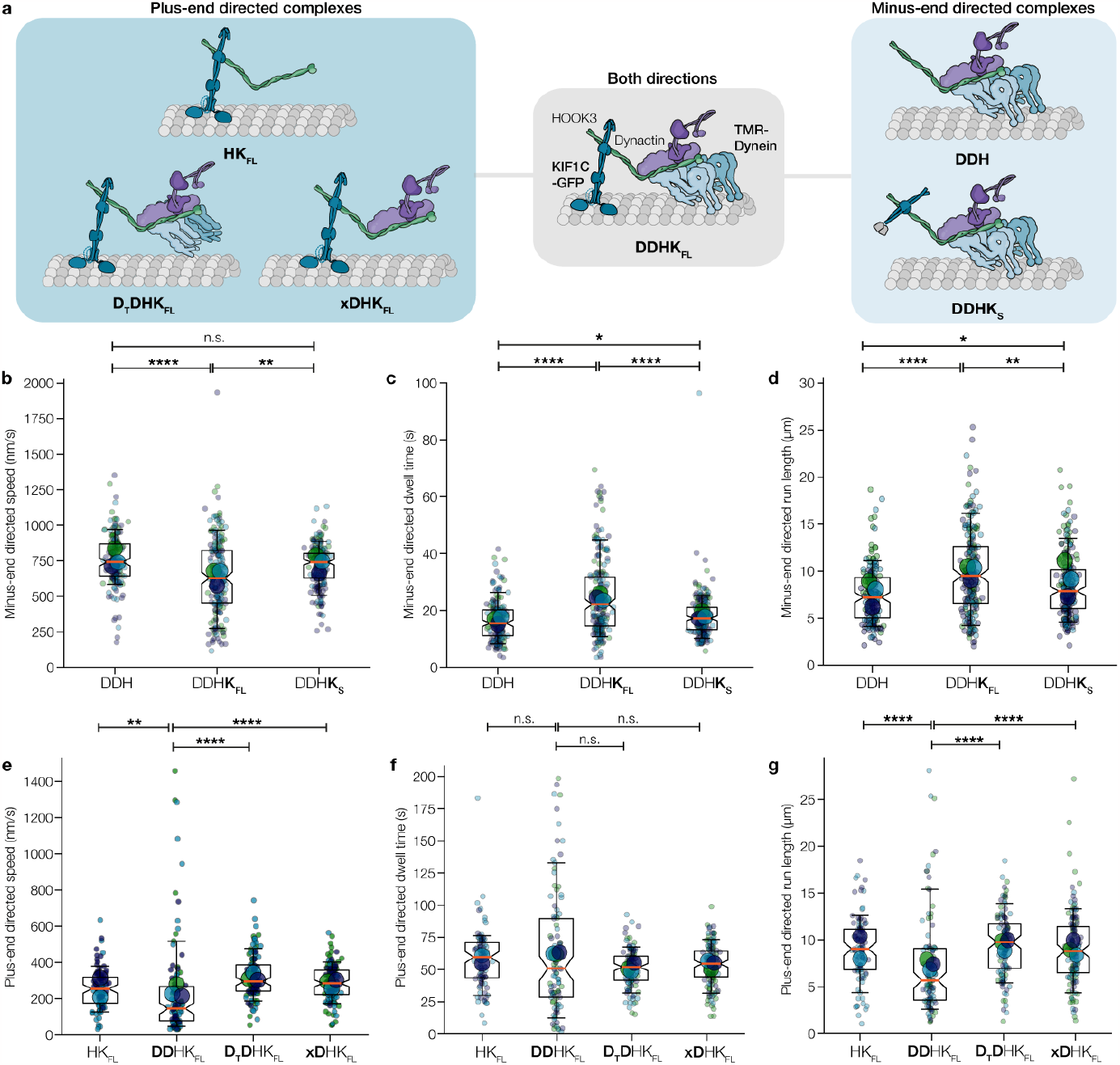
Motility behaviour of dynein and kinesin-driven co-complexes. (a) Cartoons depicting the proteins used in each co-complex. (b-d) Superplots showing minus-end directed speed, dwell-time and run lengths of dynein-dynactin-HOOK3 complexes (DDH) in the presence of either KIF1C full-length (DDHKFL) or KIF1C stalk (DDHKS). Small dots show data per microtubule analysed, large dots show experimental averages. Boxes show quartiles, whiskers 10-90% of data. (e-g) Superplots showing plus-end directed speed, dwell-time and run lengths of KIF1C-HOOK3 complexes (HKFL) in the presence of dynein and dynactin (DDHKFL), dynein tail and dynactin (DTDHKFL) and dynactin only (xDHKFL). Small dots show data per microtubule analysed, large dots show experimental averages. Boxes show quartiles, whiskers 10-90% of data. n=94-201 microtubules. n.s. p>0.05, *p<0.05, **p<0.01, ***p<0.001, ****p<0.0001 using Kruskal Wallis H test followed by a Conover’s posthoc test to evaluate pairwise interactions with a multiple comparison correction applied using Holm–Bonferroni.

Analysis of kinesin driven motility indicated that the duration of plus end-directed runs was unchanged, but both speed and run length decreased moderately when the dynein motor domain was present (Fig. 4e-g). This suggests dynein is engaged with the microtubule during KIF1C-driven runs. However, despite interacting with the microtubules, neither KIF1C nor dynein cause stalling during opposite polarity runs when they are coupled via HOOK3 (Extended Data Fig. 9). The increased processivity of dynein in the presence of KIF1C is another mechanism by which motor co-dependence occurs.

## DISCUSSION

### A general mechanism for co-dependence?

Previously, several models were proposed to explain co-dependence of opposite polarity motors: mechanical activation, steric disinhibition and microtubule tethering^2^. Our data provide evidence for two of those mechanisms occurring sequentially: Firstly, dynein and kinesin relieve the inhibited state of their opposing counterpart to increase landing rates on microtubules (steric disinhibition model). Once dynein-mediated transport is initiated, KIF1C remains in a weak microtubule binding state, preventing detachment of cargo from the microtubule (microtubule tethering model).

The initially proposed steric disinhibition model envisioned that both motors directly interact^2^, here we propose that disinhibition instead occurs through regulating the scaffolding adaptor HOOK3. This explains why the motor domains of KIF1C and dynein are dispensable for reciprocal activation (Fig. 1, Fig. 2d-e). Although not yet formally tested, our described mechanism of motor co-dependence may be applicable to other adaptor-scaffolded ensembles. For example, KIF1C has also been shown to bind dynein via BICDR1^20^ or BICD2^42^. These adaptors recruit dynein via their N-termini^16,21,43^ and are thought to bind KIF1C at their C-termini^20,42^, akin to HOOK3 co-complexes. More recently, studies in filamentous fungi suggest that auto-inhibited kinesin-1 facilitates HookA-mediated dynein transport of endosomal cargo^25^. It is compelling to compare these examples to adaptors where dynein and kinesin bind at overlapping sites, such as for TRAK^27^ or JIP3^30^. In these cases, do motors still reciprocally activate through the adaptor?

We focused on KIF1C’s activation of dynein as the majority of the movement we observed was minus-end directed and because the network of interactions between KIF1C and FHF were previously elusive. However, we suggest activation of KIF1C movement by dynein may occur in the same way. Previous structural work showed that the HOOK3 N-terminus runs the length of dynactin and contacts the dynein light intermediate chains^21,43,44^. We also showed that the pointed end of dynactin alone can bind the adaptor^45^. At sufficiently high local concentrations of dynein-dynactin, these contacts could alleviate HOOK3 inhibition to expose the C-terminus for KIF1C binding. Indeed, our structural modelling suggests that dynein-dynactin binding would disrupt the intramolecular interactions in FHF and we observed higher KIF1C-HOOK3 recruitment to microtubules both in the presence of the full and partial dynein machinery (Fig. 1f-g).

What is the benefit of reciprocal motor activation? The pre-requisite of having kinesin activate dynein movement (and vice versa) ensures the presence of both motors on cargo before effective transport can occur. This also means that the non-driving motor, in its weak microtubule binding state, is available to act as a processivity factor. In addition, the non-driving motor could respond to cellular cues and obstacles to trigger a directional switch in a timely manner. Finally, ensuring both motors are present would also be an efficient mechanism for motor recycling so that they are available at their respective start points^46–50^.

### FHF is an autoinhibited adaptor to ensure regulated dual-motor binding

Our cryo-EM structure of FHF revealed a complex that is held together by a small region of the HOOK3 C-terminus. While all HOOK family members (HOOK1-3) bind FTS, there is evidence they show preference for different FHIP isoforms (FHIP1A, 1B, 2A and 2B)^34^. The universal nature of the FTS interaction is due to a highly conserved EEKxxxSAWYN motif found in all HOOK proteins (Extended Data Fig. 10a). This motif is present on both sides of the coiled-coil and interacts with FTS at sites 2 and 3. Consistent with this motif being important, mutations in it abolish HOOK1 binding to FTS^35^.

Site 1 forms the only point of contact between HOOK3 and FHIP1B. Surprisingly, the CTE helix, found at this site, is also highly conserved between HOOK family members (Extended Data Fig. 10a). This suggests that HOOK-FHIP specificity is either mediated by the preceding CTE loop, which varies in sequence and length among HOOKs, or by another mechanism such as specific co-translation of selected mRNAs^51^. To test the former model, we co-expressed HOOK2 or HOOK3 with different combinations of their preferred FHF partners (FTS+FHIP2A and FTS+FHIP1B respectively) (Extended Data Fig. 10b)^34^. We found all combinations produced stable FHF complexes (Extended Data Fig. 10c-d) suggesting there can be only subtle specificity based on sequence alone. Thus, we favour the hypothesis that FHF specificity is driven by a cell-based mechanism.

Other adaptors use their C-terminus to directly bind small G-proteins which tether them to their membrane cargos and activate them^52–54^. HOOK3, in contrast, is likely to bind its small G-protein (Rab5) via a site on FHIP1B that is well separated from the coiled-coil. This suggests FHF is recruited to its endosome cargo^24,25,36,55^ in its inhibited conformation (Fig. 5a, step 1). Once bound to cargo, we propose FHF binds KIF1C’s stalk (step 2), leading to a partial opening of HOOK3 that allows dynein/dynactin recruitment (step 3). As the KIF1C stalk does not induce DDFHF motility to the same level as the truncated HOOK3 control, it is possible that other, as yet unidentified, factors are required for full adaptor opening and activation.

**Figure 5:**
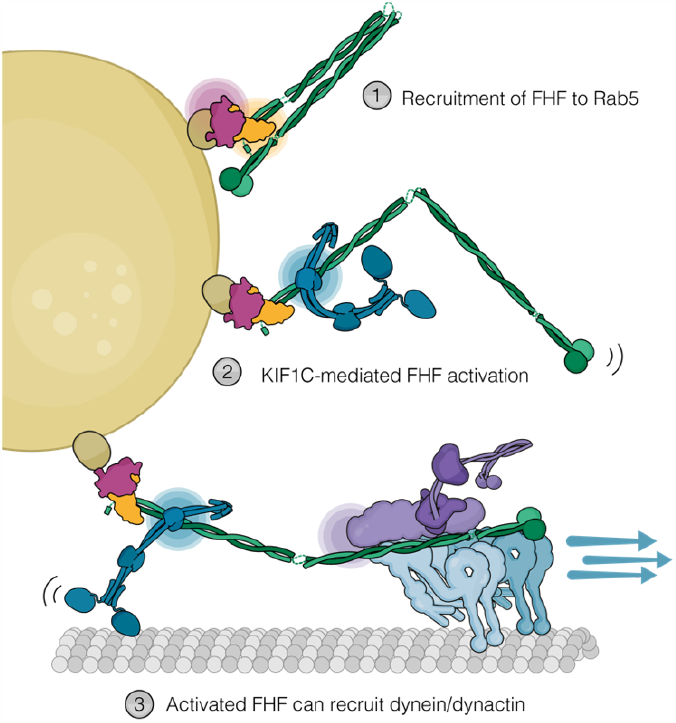
Proposed model for activation of dynein movement by KIF1C. (1) FHF assembles on Rab5 marked early endosomes. The FHF complex is inherently auto-inhibited through HOOK3 intramolecular interactions. (2) KIF1C binding (through its stalk domain) to HOOK3 C-terminus relieves HOOK3 fold-back conformation, thereby enabling (3) dynein-dynactin access to the HOOK3 N-terminus and processive minus end-motility. During these runs, KIF1C only partially engages the microtubules, acting as a processivity factor for dynein.

Our data suggest that KIF1C is partially engaged with the microtubules during dynein-driven movement (Fig. 5a, step 3). This explains how the presence of KIF1C can reduce dynein’s velocity but not stall it as it would if it were fully engaged. Our previous optical trapping experiments showed that KIF1C tends to slip backwards under load rather than fully detach from the microtubule^56^. This would potentially allow KIF1C to act as a processivity factor. However, it is unlikely that the biophysical properties of the motor alone is sufficient to explain the absence of tug-of-war in DDHK complexes, because the related kinesin-3 KIF1A has an even higher propensity for backslipping^56^, but when truncated KIF1A was coupled to dynein using a DNA-based linker, the majority of complexes showed little net velocity^33^. Comparing the plus end-directed movement of DDHK complexes containing full-length dynein with those assembled with the dynein tail (DTDHK), shows a moderate reduction in speed (Fig. 4e). This indicates that dynein is also in a conformation that causes resistance but does not stall plus-end movement. A fully engaged dynein-dynactin-HOOK3 complex has an average stall force of about 5 pN^21^, whereas only 1 pN resisting force is needed to reduce KIF1C velocity by half^56^. Therefore, during plus end-directed runs of DDHK complexes, dynein engages only weakly with microtubules to keep the resisting force below 1 pN, which is about half of the stall force of a dynein dimer^31^.

Several recent studies on adaptor-mediated dual-motor complexes also showed motility^22,27,57^. This suggests that the linkage between dynein and the opposing kinesin is crucial for the synergy of the motors. Therefore, the compliance of the cargo-adaptor-mediated linkage, steric constraints through dynactin, additional interactions or conformational changes within the DDHK complex might enable the cooperation of KIF1C and dynein. Our reconstituted system and structural insights into the engagement of both motors with the adaptor will make it possible to start to reveal these mechanisms.

## METHODS

### Cloning of protein expression constructs

Insect expression vector pFastBacM13-8xHis-ZZ-LTLT-HOOK3-SNAPf was made by amplifying HOOK3 from a human cDNA library by PCR using oligonucleotides AS689 (ACATAggcgcgccctATGTTCAGCGTAGAGTCGCT G) which encoded a 5’ AscI site and AS691 (ACAATaaccggtggatcCCTTGCTGTGGCCGGCTG), which introduced a 3’ AgeI/BamHI site. PCR products were cut with AscI and AgeI/BamHI and ligated into pFastBac-M13-8xHis-ZZ-LTLT-BICDR1-SNAPf opened with AscI and BamHI/AgeI sites to replacing BICDR1. This attached an N-terminal purification tag containing 8xHis, tandem protein-A binding motifs (ZZ) and tandem TEV recognition sites (LTLT) and a C-terminal SNAPf domain.

An expression vector for HOOK3 without a C-terminal tag was produced by PCR using AS689 and AS967 (ATAAATgcggccgcCGaccggtttaCCTTGCTGTGGCCGGCTG). This was cut with AscI and NotI and then inserted into pFastBac-M13-8xHis-ZZ-LTLT-HOOK3-SNAPf replacing HOOK3-SNAPf with HOOK3 followed by a stop codon and creating pFastBac-M13-8xHis-ZZ-LTLT-HOOK3.

For all FHF plasmids used in this study, the three human FHF genes were first codon optimised for expression in Sf9 cells and synthesised commercially by Twist or Epoch Life Science. The genes were first subcloned into pACEBac1 vectors using Gibson cloning^58^. In the case of HOOK, the full-length or truncated form of the gene (e.g. H^N^ or H^c^) was subcloned into a pACEBac1 vector encoding a Strep II tag followed by a Prescission protease cleavage site (added in frame at the 5’ end). A pBig1 plasmid for FHF co-expression was assembled using the biGBac system as described previously^40,59^, where the pACEBac1 vectors were used as backbone. Correct assembly of FHF was verified by Sanger sequencing (Source Bioscience). Constitutively active HOOK3^1-522^ was constructed as described previously^21^ .

The GST-KIF1C stalk construct was cloned into a modified *pGEX-6P2* vector (*pGEX-6P2-LTLT*) which contains an insertion of a tandem TEV cleavage site between the *BamH*I and *Eco*RI sites. Amino acids 642 to 922 from human KIF1C were amplified from *pKIF1C-2xFlag* using AS696 (GAGTCgaattcGAAATGGAGAAGAGGCTGCAG) and AS778 (ATAAATgcggccgcCTACTCCCAGCTTGACAGTGG TG), digested with *Eco*RI and *Not*I, and ligated into *pGEX-6P2-LTLT* to create *pGEX-6P2-LTLT-KIF1C(642–922)*. The GST-KIF1C stalk-GFP (pET22b-GST-6P2-LTLT-KIF1C(642–922)-GFP) bacterial expression plasmid was created by opening pET22b-HsCLASP2-EBBD-eGFP6-His between the NdeI and SalI sites, and ligating in a PCR product of GST-KIF1C stalk (previously described) formed between oligonucleotides AS777 (TGCCAcatATGTCCCCTATACTAGGTTATT) and AS780 (ATATTgtcgaccCTCCCAGCTTGACAG).

All other variants of GST-KIF1C stalk construct (KIF1CS^2^: 674-922, KIF1CS^3^: 674-822, KIF1CS H3BD: 720-841, KIF1C stalk ∆H3BD_1: 642-922 ∆722-840, KIF1C stalk ∆H3BD_2: 642-922 ∆722-840) were based on the original GST-KIF1C stalk plasmid, where fragments were PCR amplified at the indicated sequence positions and assembled using Gibson cloning^58^.

### Expression of Hook3

Sf9 insect cells (VWR, #EM71104-3) were maintained in ExCell 420 serum-free media (Sigma, 14420C). For baculovirus expression of labelled SNAP-tagged adaptor proteins, pFastBac-M13-derived plasmids were transformed into DH10BacYFP competent cells and plated on LB-Agar supplemented with 30 μg/ml kanamycin (#K4000, Sigma), 7 μg/ml gentamycin (#G1372, Sigma), 10 μg/ml tetracycline (#T3258, Sigma, 40 μg/ml Isopropyl β-D-1-thiogalactopyranoside (IPTG, #MB1008, Melford) and 100 μg/ml X-Gal (#MB1001, Melford). Positive transformants (white colonies) were screened by PCR using M13 forward and reverse primers for the integration into the viral genome. The bacmid DNA was isolated from the positive transformants by the alkaline lysis method and transfected into SF9 cells with Escort IV (#L-3287, Sigma) according to the manufacturer’s protocols. After 5–7 days, the virus (passage 1, P1) is harvested by centrifugation at 300 × g for 5 min in a swing out 5804 S-4-72 rotor (Eppendorf). P1 virus was used to infect 50 ml of SF9 culture and P2 virus was harvested after 3-5 days through centrifugation and subsequent filtration through 0.45 μm PVDF membrane (Merck Millipore, SLHV033RS). P2 virus was either propagated further or stored at 4ºC in the dark. For protein expression, Sf9 cells were infected at a density between 1.52x10^6^ cells/ml with 1:100 dilution of P2 or P3 virus. Cultures grown for 48-72 hours or until all cells contained YFP fluorescence under the microscope. Cells were harvested by centrifugation at 500 × g for 15 minutes, and pellets were stored at 80°C until used.

### Expression of FHF

All FTS-Hook-FHIP constructs were expressed using the baculovirus/Sf9 system. Bacmid DNA was generated by transformation into DH10EmBacY cells (comprising a YFP reporter gene). White colonies were picked from selective plates and bacmid DNA were recovered using a modified Qiagen mini-prep protocol. To generate P1 virus, 2-3 µg bacmid DNA was transfected (FuGene HD, Promega) into 2 ml Sf9 cells plated at 0.5x10^6^ cells/ml density in a 6-well plate and left to incubate for 5-7 days 27ºC without shaking. Success of transfection was monitored by the intensity of green fluorescence emitted from cells under a fluorescence microscope. These P1 virus were harvested by pipetting and amplified to create P2 virus: 1 ml P1 was added to 50 ml Sf9 cells at 0.5x10^6^ cells/ml in a 250 ml Erlenmeyer flask to give a 1:50 v/v dilution. This was incubated at 27 ºC, 140 rpm shaking for 72 hours. P2 virus were harvested (supernatant) by spinning at 3750 rpm in a Beckman centrifuge for 5 min at room temperature. For protein expression, 5-7 ml of P2 virus was added to 500 ml of Sf9 cells at 1.5x10^6^ cells/ml in roller bottles and left to incubate at 27 ºC, 140 rpm shaking for 56 hours. At this time point the cells were harvested by centrifugation (4,000 rcf for 10 min at 4 ºC) and snap frozen in liquid nitrogen prior to storage at -80 ºC.

### Purification of full-length HOOK3

Unless otherwise stated, all protein purification procedures were performed at 4°C. SF9 cells were lysed in lysis buffer (50mM HEPES pH 7.4, 150mM NaCl, 20mM Imidazole) supplemented with 1x cOmplete protease inhibitor cocktail (Roche, 6538282001). Lysis was achieved in a glass tissue douncer with the tight pestle using 20-30 strokes. The resultant lysate was cleared by centrifugation at 50,000xg for 40 minutes. Cleared lysate was loaded on to 12ml of NiNTA agarose beads (Qiagen, 30250) and batchbound for 1.5 hours. Beads were transferred to a gravity flow column and washed with 100-200 CVs of lysis buffer followed by 100-200 CVs of wash buffer (50mM HEPES pH 7.4, 150mM NaCl, 60mM Imidazole). Protein was eluted in up to 10ml of elution buffer (50mM HEPES, pH 7.4, 150mM NaCl, 300mM Imidazole), which was subsequently batchbound to 1ml IgG Sepharose (Cytiva, 17096901) for 1.5 hours. Proteinbound IgG beads were washed with 100 CVs of TEV Cleavage buffer (50mM Tris pH 7.4, 148mM KAc, 2mM MgAc, 1mM EGTA, 10% Glycerol), transferred to a 1.5ml tube and incubated with 3-5 μM benzylguanine conjugated fluorophore (NEB, S9136S) for 2 hours before being returned to the column, washed with a further 100 CVs of TEV Cleavage buffer, and cut off the beads with 40 μg/ml TEV protease at 25 °C for 1 hour. The protein was collected from the beads and the bead bed was washed with a further 1 CV of buffer. The collected protein was subjected to a clearing spin at 13,000 rpm at 4 °C for 5 minutes before being aliquoted and stored in N2 vapour.

Untagged HOOK3 was expressed from pFastBac-M13-8xHis-ZZ-LTLT-HOOK3 as described above and purified by a singlestep IgG sepharose purification followed by gel filtration over a Superose 6 increase 10/300 column. The purification buffer used throughout was 50 mM HEPES 7.5, 150 mM NaCl, 2.5 mM MgSO4, 10% Glycerol, 2 mM DTT, 0.1 mM ATP and 0.05% Triton X. Lysis was performed using 20-30 strokes in tissue douncer, cleared lysates were bound to 750 μl of IgG sepharose beads for 3 hours, and the beads were washed with 300-400 ml of purification buffer. Beads were resuspended in a 1.5 ml tube and incubated with 40 μg/ml TEV protease for 1 hour at 25 °C with agitation. Eluate was collected in a gravity flow column, the beads were washed with an additional 1 CV of buffer, and then the protein was concentrated using a 30 KDa centrifugal concentrator to a final volume of around 600 μl. The protein was clear spun at 13,000 rpm at 4 °C for 5 minutes before being injected into a pre-equilibrated Superose 6 increase 10/300 column in gel filtration buffer (50 mM HEPES 7.4, 150 mM NaCl, 2.5mM MgSO4, 1 mM DTT). The peak corresponding to HOOK3 at around 11-12 ml was concentrated in the presence of 10% glycerol and then aliquotted, snapfrozen, and stored in N_2_ vapour.

### Purification of FHF complexes

2L worth of pellets were thawed in a water bath and re-suspended with 45 ml lysis buffer (50 mM Hepes pH 7.2, 150 mM NaCl, 10% glycerol, 1 mM DTT) supplemented with 2 mM PMSF and two complete-EDTA free protease inhibitor tablets per 50 ml of buffer. Cells were lysed using a tight Dounce homogenizer using 20-25 strokes on ice. Cell debris was pelleted by spinning lysates at 70,000 rpm for 45 min at 4ºC in a Ti70 in an ultracentrifuge (Beckman Coulter). In a cold room, the clarified lysate was applied to 4 ml pre-equilibrated Strep-Tactin beads (lysis buffer) in a Bio-Rad gravity flow column and the flow-through was immediately collected. The flow-through was re-applied to the beads to ensure full capture of protein. Beads were washed with ~450 ml lysis buffer followed by 200 ml Psc buffer (50 mM Tris-HCl pH 7.4, 150 mM NaCl, 1 mM EDTA, 1 mM DTT). Beads + 6 ml Psc buffer were collected with cut pipette tip and distributed equally across two 5 ml Eppendorf tubes. To cleave bound protein from beads, 90 µl Psc enzyme at 2 mg/ml was added to each tube and left to incubate in a PTR-60 Multi-Rotator (Grant-bio) at orbital setting 2 for 16 hours at 4°C. The following day, the bead-elution mixture was added to a small gravity column and washed with 5 ml Psc buffer to release remaining loosely bound protein. The elution was concentrated in a 15 ml 100kD cut-off (4,000 rpm, 5 min spins) and mixed in between spins by pipetting until reaching ~8 mg/ml concentration in ~800 µl volume. Concentrated protein was spun down in a microfuge tube within a microcentrifuge (12,000 rcf for 5 min). 250 µl protein was injected into a Superose 6 10/300 column using a 500 µl loop, collecting 100 µl fractions. Peak fractions were pooled and concentrated in a 4 ml 100kD-cut off concentrator. 10% glycerol was added, and protein was aliquoted into 5 µl lots. Typical yields from these purifications ranged from 2-4 mg.

### Purification of KIF1C stalk constructs

All KIF1C stalk constructs used in this study, apart from GST-KIF1C stalk-GFP, were purified as follows. 1-2 L worth of SoluBL21 pellets expressing the relevant GST KIF1C stalk construct were resuspended to a homogenous mixture with 50 ml lysis buffer (50 mM Tris pH 7.4, 300 mM NaCl, 20 mM imidazole, 10% glycerol, 1 mM DTT) supplemented with 2 mM PMSF + cOmplete™, EDTA-free Protease Inhibitor Cocktail tablet. Cells were lysed by sonication and lysate was cleared by spinning down in a Ti70 at 65K rpm (4ºC) for 30 min (Beckman Coulter). Cleared lysate was briefly incubated (approximately 5 min) with 3 ml Ni-NTA Agarose (QIAGEN) before flow-through was collected. Beads were washed with approximately 250 ml lysis buffer followed by 200 ml high salt lysis buffer (50 mM Tris pH 7.4, 500 mM NaCl, 30 mM imidazole, 10% glycerol, 1 mM DTT) and back to an additional 200 ml lysis buffer. Eluted protein using elution buffer (50 mM Tris pH 7.4, 300 mM NaCl, 300 mM imidazole, 1 mM DTT) in 1 ml increments until 15 ml was collected. Concentrated down combined elution in a 30-100K cut-off 15 ml concentrator (depending on KIF1C stalk construct size) prior to injection into a Superdex 200 increase 10/300 column (Cytvia) pre-equilibrated in GF300 (25 mM HEPES 7.2, 300 mM NaCl, 1 mM MgCl2, 1 mM DTT.

Peak fractions were pooled from multiple consecutive runs and concentrated down in a 30K cut-off 4 ml concentrator until ~150-200 µl at 3 mg/ml. Fractions were snap frozen and stored at - 80ºC until further use.

### Purification of other proteins

KIF1C-GFP and GST-KIF1Cstalk-GFP constructs^23^, dynactin^21^, TMR-labelled human dynein^15^ and Lis1^60^ were expressed and purified as described previously.

### TIRF assays of HOOK3-containing motor complexes

Menzel Gläser No. 1.5 22x22 mm and 22x50mm coverslips were placed in holding racks and submerged in 6.4% wt/v HCl at 60 °C for 16 hours. The HCl was exchanged for distilled water, and the coverslips were sonicated in a ultrasonic waterbath for 5 minutes at a time, exchanging the distilled water between sonications. The sonication step was repeated 5 times and then coverslips were dried with compressed air. Coverslips were stored between layers of lens tissue (Ross Optical, AG806) inside pipette boxes until they were used. Prior to imaging chamber assembly, the acidwashed coverslips were plasma cleaned in a Henniker plasma clean (Henniker Plasma, HPT200) for 5 minutes. For each flow chamber, a 75x25mm SuperFrost Plus glass slide (Thermo Scientific, J1800AMNZ) was taken and doublesided tape (Tesa, 64621) was laid in two parallel lines with a gap of 5mm in between them. A single coverslip was laid over the tape and flattened to create an enclosed chamber with a volume of approximately 10-15 μl. In experiments where additional components would be added to the TIRF flowchamber while it was on the microscope, an open chamber design was adopted. Chambers were constructed between a 22x50 mm coverslip with a 22x22 mm coverslip on top. By imaging through the 22x50 mm coverslip, the top of the chamber was left open allowing additional components to be flowed into the chamber in situ. These chambers were not as hydrophilic as those created with the SuperFrost Plus coated glass slides, and therefore they were prewet with 0.1% v/v Triton X100 to prevent formation of air gaps during filling.

Microtubules were polymerised in a final volume of 20 μl MRB80 (80mM PIPES pH 6.8, 4mM MgCl2, 1mM EGTA, 1mM DTT) using 82 μg of unlabelled porcine tubulin with 2.5 μg of biotinlabelled porcine tubulin (Cytoskeleton, T333PB). The mixture of diluted tubulin was pipetted to mix, and then spun for 5 minutes in a chilled airfuge (Beckman Coulter, 340401) at around 80,000 rpm. The spun tubulin mixture was placed into a 0.5 ml tube, a final concentration of 5 mM GTP was added, and microtubules were allowed to polymerise for 1 hour at 37 °C. After 1 hour, the mixture was topped up to 100 μl by the addition of 80 μl MRB80 containing 50 μM paclitaxel (Alfa Aesar, J62734.MC), flicked to mix, and kept at room temperature. The next day, microtubules were pelleted in a benchtop centrifuge at 13,000 rpm for 12 minutes, the supernatant containing any unpolymerised tubulin was removed and replaced with 100 μl MRB80 containing 50 μM paclitaxel. The microtubule pellet was flicked to resuspend it, and once it had been broken up, the microtubules were pipetted gently 5-10 times with a 200 μl pipette tip until no traces of the pellet were remaining. Microtubules prepared in this way were kept for 3-4 weeks with the tube wrapped in aluminium foil, and diluted 1:50 in taxolcontaining buffer prior to use in microscopy chambers.

Throughout the following steps, flow chambers were kept upturned in humidified chambers to avoid drying. Chambers were first coated with PLL(20)g[3.5]PEG(2)/PEG(3.4) biotin(50%) (Susos) by flowing in 12 μl of a 0.2 mg/ml solution in MRB80. Chambers were incubated for at least 10 minutes before being washed with 30μl TIRF assay buffer (TAB: 25 mM HEPES pH 7.2, 5 mM MgSO4, 1mM EGTA, 1mM DTT, 10 μM paclitaxel), and then 12 μl of 0.625 mg/ml streptavidin (Merck, S4762) in TAB was added. Excess streptavidin was washed away with 30μl TAB before 14 μl of 1:50 diluted polymerised GDPtaxol microtubules were added to the chamber. The microtubules were allowed to attach for 10-20 seconds before unstuck microtubules were washed away with 14 μl TAB. The chamber was blocked for a period of 1-2 minutes using 1 mg/ml κcasein (Merck, C0406) in TAB buffer. Excess κcasein was washed away with 14 μl TAB and then reaction mixtures were flowed into the chamber.

The reaction mixtures were prepared in TAB supplemented with 0.2 mg/ml κcasein, 25 mM KCl, 20 μM paclitaxel, an ATP regenerating system (5 mM ATP (Melford, B3003), 5 mM phosphocreatine (Merck, P7936), 7 U/ml creatine phosphokinase (Merck, C3755)) as well as an oxygen scavenger system (0.2 mg/ml catalase (Merck, C9322), 0.4 mg/ml glucose oxidase (Merck, G7141), 4 mM DTT and 50 mM glucose). 200 nM Dynein, 200 nM dynactin, 1600 nM labelled adaptors and 180 nM KIF1CGFP were mixed in 2.5 µl of dynein storage buffer GF150 (25 mM HEPES 7.4, 150 mM KCl) and then diluted with 2.5µl TAB buffer which contains no added salt, and thus the final salt concentration during complex assembly was ~75 mM KCl. As the concentration of untagged hook3 was lower than that of tagged cargo adaptors, an excess of Lis1 was added to aid complex formation. Complexes with unlabelled HOOK3 were formed by mixing 50 nM of each dynein, dynactin and KIF1CGFP or KIF1C buffer control in the presence of 250nM untagged HOOK3 and 1250nM untagged Lis1. The complex mixture was pipetted, spun down, and then retained on ice for 45-60 minutes. Just before use, the complexes were mixed once more by pipetting before being diluted 1:20 or 1:40 into the reaction mix. The reaction mix was added to the flow chamber and imaging started immediately using an Olympus TIRF system with a ×100 NA 1.49 objective, 488, 561 and 640 nm laser lines, an ImageEM emCCD camera (Hamamatsu Photonics) under the control of xCellence software (Olympus), an environmental chamber maintained at 25 °C (Okolab, Ottaviano, Italy). Emission resulting from illumination with 488 or 561 nm laser lines was filtered by bandpass filters controlled by a filter wheel (Olympus) to avoid bleed-through.

### Motility analysis of HOOK3-containing motor complexes

Microtubules were manually traced in a maximum intensity projection or reference image, and then kymographs were generated for all the channels that were recorded using a custom ImageJ macro. ROIs were saved to return to the microtubule path if required. Next, motor tracks were manually traced with multisegmented lines in kymographs and the phase durations and lengths and speed were stored in a .csv file, while the line was also saved as region of interest (ROI) file. A python analysis software package reads the saved tracks and classifies and summarised the motility parameters (run length, dwell times, run speeds, landing rates and directionality) at the single motor level, the permicrotubule level, the permovie level and the perexperiment level. To do this, paths were segmented into runs. A run was defined as the distance covered before falling static (having an absolute speed of less than 25 nm/s), or the distance covered before changing direction (having a speed of more than 25 nm/s in one direction and then a speed of more than 25 nm/s in the opposite direction). For each segment of the path, it was determined whether this segment was going towards the plus-or minus-end of the microtubule, or whether the motor was paused, and the time spent in each mode of transport was tallied. The total time a motor spend on the microtubule was its dwell time. The total run length in the plus-or minus-end direction was also tallied for the individual runs within the track. The total run length was defined as the sum of the absolute values of all individual runs for one motor. The average speed of a particle was defined as the total run length divided by the total dwell time. The landing rate was calculated as the number of tracks per kymograph width (in nm) and length (in s) and converted to µm•min^-1^ by multiplying with 60,000. The overall directionality of a track was defined in the following way. Firstly, any track with a total runlength of less than 1000 nm was classified as “static”. Any remaining track that travelled only towards the plusend was classified as “plus end-directed”, while any track that travelled only towards the minusend was classified as “minus end-directed”. Any track that travelled more than 1,500 nm in the plus and minus end directions was classified as “bidirectional”. After this classification, any remaining track that travelled more towards the plusthan the minusend was labelled “plus end-directed”, while the opposite was labelled “ minus end-directed”. Therefore a track that travelled 20,000 nm in the minusend direction and 200 nm in the plus end direction would still be classified as “minus end-directed”.

### Statistical analysis of motility of HOOK3-containing complexes

Statistical analyses were performed in Python with use of the following modules: scipy^61^ and scikit-posthocs. All statistical analyses were performed as follows. The data were tested to see if they followed a normal distribution using D’Agostino and Pearson’s test. If data were normally distributed, pairwise interactions were tested using a twotailed ttest for independent samples, and the resulting pvalues were corrected for multiple comparisons if necessary. Where ttests have been used, this is indicated in the figure legend; otherwise all statistical analyses relate to the following nonparametric testing process. If one or more experimental groups were not normally distributed, a KruskalWallis H test was used to determine if any of the medians of the experimental groups differed. If the KruskalWallis H test showed that one group was significantly different, then pairwise interactions were tested using Conover’s posthoc test. The pvalues of these pairwise interactions were corrected for multiple comparisons.

Data were plotted using matplotlib^62^, and superplots were created with a custom-made extension to matplotlib. Graphs were edited in Adobe Illustrator (Adobe), where modifications were limited to colours, line widths, spacing and text size. Microscopy images were prepared in FiJi^63^. Image manipulations were limited to scaling of contrast (between minimum and maximum values),and displaying images with different colour lookup tables. Where images from the same experiment are displayed alongside one another, their contrast is scaled equally.

### SEC reconstitution of complexes

All gel filtration reconstitution experiments were performed using a Superose 6 Increase 3.2/300 column (Cytvia) using a 100 µl loop and 50 µl loaded sample in gel filtration buffer 150 (GF150, 25 mM HEPES pH 7.2, 150 mM KCl, 1 mM MgCl2 and 1 mM DTT). This column was either connected to an ÄKTAmicro or an ÄKTApure micro system (Cytvia). A typical reconstitution used 5 µM FHF construct and 10 µM KIF1C stalk construct diluted in GF150 to bring total input sample volume to 60 µl.

### Pulldown assays

All steps were performed at room temperature. 200 µl Strep-tactin®Sepharose® (iba) slurry was added to disposable Poly-Prep® chromatography columns (Bio-Rad) for each experimental condition. Bead storage solution was allowed to flow through followed by a 2 ml wash with GF50 buffer (GF150, 25 mM HEPES pH 7.2, 50 mM KCl, 1 mM MgCl2 and 1 mM DTT). Samples comprised 5 µM streptagged bait (FH^C^F) and 10 µM H^N^ and/or 5 µM GST-tagged KIF1Cstalk construct ± HBD. All samples were diluted with GF50 to bring to 60 µl and then incubated for 15 min. 10 µl sample input was taken away and 50 µl samples loaded onto the beads to let bind for 15 min. Beads were washed twice with 1 ml of GF50. 500 µl GF50 was then added and beads were resuspended. 400 µl of this wash+bead sample was transferred to fresh microcentrifuge tubes. Beads were spun down 1000xg for 1 min and 250 µl of the supernatant was removed. To the remaining bead+buffer mixture (~150 µl volume), 30 µl of NuPage™ LDS Sample Buffer [4X] (Invitrogen) was added. 30 µl of this bead sample was loaded into a 10-well NuPAGE™ 4 to 12%, Bis-Tris, 1.5 mm, Mini Protein Gel and ran using MOPS buffer.

### Mass Photometry

To assess the oligomeric state of FHF with and without KIF1C stalk, we used a mass photometry machine (TwoMP version, Refeyn Ltd). ~10 µl samples were added to wells within gaskets attached to clean coverslips. These samples were diluted to 100 nM in GF150 buffer. Movies were imaged for 60s using the AcquireMP software (Refeyn Ltd) and the results were processed using the DiscoverMP software. Gaussian fitting was used to determine the mass peaks against a calibration performed using similar buffer conditions.

### Cryo-EM sample preparation

4 µl of FHF at 0.9 mg/ml containing either 0.1% CHAPSO or 0.005% Igepal was added to an unsupported Quantifoil (2/2 Au 300-mesh) grid and left to incubate for 15-20s inside a Vitrobot mark IV prior to blotting for 1s (-13 blot force) and plunge freezing into liquid ethane. Grids were stored in liquid nitrogen and screened for ice thickness and protein concentration using a ThermoFisher Glacios TEM equipped with a Falcon III (ThermoFisher).

### Cryo-EM data collection

Electron micrograph movies were collected using a Titan Krios (300keV X-FEG) equipped with a K3 direct electron detector and a Gatan BioQuantum energy-filter (dataset 1 and 3) or Falcon 4i pre-GIF direct electron detector (dataset 2). Movies were collected at 75,000x magnification in EFTEM mode, yielding a pixel size of 1.09 Å for dataset 1 and 3 (Fringe-Free Illumination) and 1.08 Å for dataset 2. Aberration-free image shift (AFIS) collection was used for data collection (5 acquisitions/hole for K3 or 4 acquisitions/hole for Falcon 4). See Supplementary table 1 for acquisition parameter details.

### Cryo-EM image processing

All cryo-EM image processing steps were performed in RELION-4.1^64^ unless otherwise stated. Movies were motion corrected using the RELION implementation. Particles were picked in crYOLO using the general model trained on low-pass filtered images^65^. These particles were extracted (bin2, giving a pixel size of 2.16-2.18 Å/pixel depending on the camera). At this point, datasets 1+2 were combined while dataset 3 was kept separate until later (see Extended Data Fig. 2b). For datasets 1+2 (total: 2,507,930 particles), particles were subjected to unmasked 3D classification (5 classes) with alignment using an ab-initio generated initial model of FHF. The best class was selected (1,012,409 particles) and a global 3D refinement was performed, yielding an overall 4.8 Å resolution structure. This was Bayesian polished and rerefined, leading to an improved 4.5 Å resolution structure. At this point, particles were re-extracted unbinned and subjected to a round of masked 3D classification without alignment. This gave two good classes, which were selected (781,492) for subsequent merging with dataset 3 particles.

Dataset 3 was processed as follows. 2,368,723 extracted particles were used for cleaning steps using 2D classification. 1,232,461 particles were taken forward for unmasked 3D classification with alignment and the best class selected (723,461 particles) was subjected to 3D refinement (using global search parameters), giving a 4.4 Å structure. Dataset 3 particles were then re-rextracted to their unbinned parameters (256 box size, 1.09 Å/pix). The unbinned particles were used for a local 3D refinement, resulting in a 3.7 Å structure. Bayesian polishing was performed using this unbinned, consensus refinement, which improved the subsequent 3D refinement to 3.3 Å resolution. However, the structure was highly anisotropic at this point. To reduce anisotropy in the map, a combination of StarParser (https://github.com/sami-chaaban/starparser) and RELION’s subset select tool was used to remove overrepresented views. StarParser was used to plot a histogram of the rlnAngleRot values from the 3D refinement particles.star file. Based on this visualisation, the rlnAngleRot values were split into different bins using RELION’s subset selection (metadata label: rlnAnglRot), e.g. angles that fill in within -140 to 100 and between 100 to 200. This generated two particles star files. The latter star file (100-200 rlnAngleRot) had the majority of overrepresented views and this was subjected to random extraction of ~50% of particles to reduce the overrepresentation. The output star file was then combined with the unmodified first star file (-140-100 rlnAngle) (Join star files in RELION) and duplicate particles were removed (Subset selection tool).

At this point, datasets 1+2 and dataset 3 particles were merged and a new consensus refinement was obtained of the merged dataset (total particles: 1,207,965) at 3.4 Å resolution. These particles were subjected to 3D classification without alignment (8 classes, T=8, with a mask around the FHF [8 pixels binary map extension and 8 pixels soft edge]). The class with the highest resolution features and most complete density was selected, giving 377,595 particles. As the overall structure still showed anisotropy at this point, the reduce anisotropy procedure described above was employed at this point. 273,204 particles were subsequently taken forward to 3D refinement, CTF refinement (defocus and astigmatism), 3D refinement, Ctf refinement (magnification anisotropy) before a final 3D refinement. After postprocessing, this final map was at 3.2 Å resolution (after correction of calibrated pixel size at 1.059 Å/pixel), and was used for all subsequent model building and real-space refinement.

### Model building and refinement

The initial coordinates for docking into the FHF cryo-EM density were predicted using a local cluster installation of Colabfold 1.5.0^66^ (AlphaFold2^67^), where full-length sequences of human FHIP1B and FTS were provided alongside a dimer of truncated HOOK3 (571-718). This resulted in a good overall fit but required dividing up of FHIP1B into two halves to better fit the N and C-termini of the alpha solenoid. In addition, a switch had to be made to place the correct loop-CTE helix-loop region within the HOOK3 density (the other CTE helix was cut out from the model as no density could be observed for this region). Coot^68^ was also used for real-space refinement of backbone and sidechains within the density. Side chains were removed from low resolution density regions using Phenix^69^ (version 1.14). The model was then real-space refined using Phenix (version 1.20) after adding hydrogens (using a reference_coordinate_restraints.sigma value of 0.0005). Hydrogens were removed and the resulting model was inspected in Coot, where outliers were fixed before repeating the real-space refinement process one last time. All structural figures were prepared using UCSF Chimera X^70^.

### AlphaFold predictions

AlphaFold structures of FHF and FHF bound to kinesin stalk were predicted using a local cluster installation of Colabfold 1.5.0^66^ where homology searches were performed using MMseqs2^71^ and Alphafold2-Multimer (v3) was utilised^67,72^. AlphaScreen (https://github.com/sami-chaaban/alphascreen) was used

### Analysis of processive events in FHF-containing samples

Microscope coverslips (epredia 22 × 22 mm # 1.5) were cleaned by loading onto ceramic holders, soaking in ~300 ml 3M KOH in a 1L beaker and sonicating for 30 minutes in a water bath sonicator. Coverslips were washed by transferring to a new beaker containing ~500 ml MQ H2O and washed thoroughly five more times using the same volume of water. Coverslips were sonicated in MQ H2O for 30 min and transferred to new beaker containing 96% ethanol. Ethanol immersed coverslips were sonicated for 30 minutes and stored in fresh ethanol. Prior to use, ethanol was removed from the coverslips using compressed nitrogen and placed in their ceramic holder in a plasma cleaner (Fischione instruments, model 1070 NanoClean) and evacuated until “High Vac” was reached. Coverslips were exposed to plasma for 3 minutes with a setup of 75% Argon and 25% Oxygen at 100% power.

Motility chambers were prepared as previously described^73^. Briefly, two strips of double-sided tape were applied to a glass slide (epredia ground 90º frosted, BS7011/2 1.0-1.2 mm) at ~5 mm apart. A pre-cleaned coverslip was placed on top, and firmly sealed, to form a channel that can take 10-15 µl of liquid.

Microtubules were prepared by adding 6 µl 13 mg/ml unlabelled pig tubulin, 2 µl 2 mg/ml Alexa fluor 647 tubulin, 2 µl 2 mg/ml biotin-tubulin in BRB80 buffer (80 mM PIPES pH 6.8, 1 mM MgCl2, 1 mM EGTA, 1 mM DTT) and 10 µl polymix buffer (2xBRB80, 20% v/v DMSO and 2 mM Mg.GTP). This mixture was left to incubate for two hours at 37 ºC in a heat block filled with water. Microtubule mixture was diluted to 100 µl with BRB80-T (BRB80 supplemented with 10 µM Taxol) and spun down at 21,300 rcf for 8.5 min at room temperature to remove excess tubulin. The pellet was washed by resuspending with 100 µl BRB80-T and flicking the bottom of the tube, then washed once more in this way. The pellet was resuspended with 80 µl BRB80-T, covered in foil and stored overnight before use. Prior to use, the microtubule pellet was resuspended by flicking and its concentration was adjusted using BRB80-T based on visualising the density of filaments inside the motility chambers.

To prepare TIRF channels, 10 µl pluronic-F127 was added to the motility chambers for 1 min prior to addition of 15 µl 0.4 mg/ml biotinylated poly(l-lysine)-g-poly(ethylene-glycol) (SuSoS AG). The channel was washed with DLB (30mM Hepes-KOH pH 7.2, 5mM MgSO4, 1mM EGTA, 1mM DTT) before adding 15 µl 1 mg/ml Streptavidin (NEB) before immediate wash with DLB-C-T buffer (DLB supplemented with 1 mg/ml α-casein, 50 mM KCl and 5 µM Taxol). A dilution was made of the microtubule stock in BRB80-T (between 1 in 3 to 1 in 5 dilution) and 15 µl of this was allowed to flow into the chamber followed by a 10 µl DLB-C-T wash. The stock complex was prepared in a 5 µl volume (diluted with GF150: 25 mM HEPES pH 7.2, 150 mM KCl, 1 mM MgCl2, 1 mM DTT) such that the molar ratios were as follows: labelled TMR(SNAPf)-dynein [100 nM]:dynactin [100 nM]:FHF [2 µM]:Lis1 [6 µM]:KIF1C-stalk [1.2 µM]. After 5 min incubation on ice, complex was diluted to 1 in 5 in buffer DLB-C-T and a further 1 in 20 in the final reaction mixture (1 µl diluted complex, 15 µl DLB-C-T, 1 µl 20 mM Mg.ATP, 1 µl catalase-glucose oxidase (0.2 mg/ml catalase [Calbiochem] mixed with 1.5 mg/ml glucose oxidase [Sigma-Aldrich]), 1 µl 9% (w/v) glucose, 1 µl 25% (v/v) 2-mercaptoethanol). 15 µl of this reaction mixture was added to the TIRF motility chamber immediately before data collection.

TIRF movies of moving molecules and snapshots of associated microtubules were collected at room temperature on a Nikon Eclipse Ti inverted microscope with a Nikon 100x TIRF 1.49 NA 100x oil immersion objective equipped with a back illuminated EMCCD camera (iXonEM+ DU‐897E, Andor, UK). µManager software was used to collect 500-frame movies at 100 ms exposure and 105 nm/pixel using a 561 nm (100 mW, Coherent Cube) laser or a 100 ms 1-frame snapshot using a 641 nm (100 mW, Coherent Cube) laser for the microtubules.

Data analysis was manually performed using FiJi^63^. Here, Z-projections of tif stacks were used to draw segmented lines along individual microtubule tracks and the reslice tool was used to make kymographs for each microtubule. Processive events for single molecules was defined as complexes that associated with microtubules for ≥1.2s and ≥525 nm (i.e. at least 5 pixels on y-axis and 5 pixels on x-axis), based on previously published criteria^15^. Velocity and run-length was calculated for each processive event using the 105 nm pixel size and 0.136 s inter-frame rate.

Statistical analysis was performed in GraphPad Prism version 9.0.0. All the individual data was plotted in column format (15 microtubules per n, n=8, therefore 120 microtubules per sample). The data were tested to see if they followed a normal distribution using D’Agostino-Person omnibus normality test, which indicated that the different experimental groups were not normally distributed. A Kruskal-Wallis H test corrected for multiple comparison (Dunn’s) was then used to test statistical significance (applied to n=120 per experimental group).

### Crosslinking mass-spectrometry

Protein cross linking reactions were carried out at room temperature for 60 minutes with 50 mg of complex present in 10 mM of EDC or 5 mM of DSSO. Crosslinked protein was quenched with the addition of Tris buffer to a final concentration of 50 mM. The quenched solution was reduced with 5 mM DTT and alkylated with 20 mM idoacetamide. An established SP3 protocol^74,75^ was used to clean-up and buffer exchange the reduced and alkylated protein, shortly; proteins are washed with ethanol using magnetic beads for protein capture and binding. The proteins were resuspended in 100 mM NH4HCO3 and were digested with trypsin (Promega, UK) at an enzyme-to-substrate ratio of 1:25, and protease max 0.1% (Promega, UK). Digestion was carried out overnight at 37 °C. Clean-up of peptide digests was carried out with HyperSep SpinTip P-20 (ThermoScientific, USA) C18 columns, using 80% Acetonitrile as the elution solvent. Peptides were then evaporated to dryness via Speed Vac. Dried peptides were suspended in 3% Acetonitrile and 0.1 % formic acid and analysed by nano-scale capillary LC-MS/MS using an Ultimate U3000 HPLC (ThermoScientific, USA) to deliver a flow of 300 nl/min. Peptides were trapped on a C18 Acclaim PepMap100 5 μm, 100 μm × 20 mm nanoViper (ThermoScientific, USA) before separation on PepMap RSLC C18, 2 μm, 100 A, 75 μm × 50 cm EasySpray column (ThermoScientific, USA). Peptides were eluted on a 90 minute gradient with acetonitrile and interfaced via an EasySpray ionisation source to a quadrupole Orbitrap mass spectrometer (Q-Exactive HFX, ThermoScientific, USA). MS data were acquired in data dependent mode with a Top-25 method, high resolution scans full mass scans were carried out ((R = 120,000, m/z 350 – 1750) followed by higher energy collision dissociation (HCD) with stepped collision energy range 21,27,33 % normalised collision energy. The tandem mass spectra were recorded (R=30,000, AGC target = 5 × 10^4^, maximum IT = 150 ms, isolation window m/z 1.6, dynamic exclusion 50 s). Cross linking data analysis: Xcalibur raw files were converted to MGF files using ProteoWizard^76^ and cross links were analysed by MeroX^77^. Searches were performed against a database containing known proteins within the complex to minimise analysis time with a decoy data base based on peptide sequence shuffling/reversing. Search conditions used 3 maximum missed cleavages with a minimum peptide length of 5, cross linking targeted residues were K, S, T, and Y, cross linking modification masses were 54.01056 Da and 85.98264 Da. Variable modifications were carbmidomethylation of cysteine (57.02146 Da) and Methionine oxidation (15.99491 Da). False discovery rate was set to 1 %, and assigned cross linked spectra were manually inspected.

## ACKNOWLEDGEMENTS

We thank S. Chaaban for setting up AlphaFold2 installations, writing and advising on AlphaScreen and StarParser code use, and advice for structural predictions. We thank C.K.Y. Lau and K. Singh for general AlphaFold2 analysis and model building advice. We thank the MRC Laboratory of Molecular Biology Electron Microscopy Facility for EM data collection time and J. Grimmett, T. Darling and I. Clayson for scientific computing support. We thank

C.M. Johnson from MRC Laboratory of Molecular Biology Biophysics facility for assistance and analysis of SEC-MALS and Mass Photometry experiments. We thank J. Shi for providing and maintaining insect cell cultures for protein expression. We thank Warwick CAMDU (Computing and Advanced Microscopy Unit) for their support & assistance. This work was supported by a Sir Henry Wellcome Postdoctoral Fellowship to F.A.A. (218653/Z/19/Z), Wellcome Investigator Awards to A.P.C (210711/Z/18/Z) and A.S. (200870/Z/16/Z & 224563/Z/21/Z), Medical Research Council funding to A.P.C (MC_UP_A025_1011) and MRC PhD studentship to A.J.Z. (MR/N014294/1).

## AUTHOR CONTRIBUTIONS

F.A.A. performed FHF-containing sample protein purification, reconstitution assays, EM data analysis, structure determination, single molecule experiments and with A.P.C. and R.F.W. cloned constructs. A.J.Z. performed the HOOK3-containing sample purifications, cloning, reconstitutions, single molecule experiments and data analysis, and with C.E.S. purified dynein and dynactin. T.E.M. with F.A.A. performed the cross-linking mass spectrometry experiments. A.S. and A.P.C. guided the project. F.A.A., A.S. and A.P.C. prepared the manuscript with input from all the authors.

---

**Extended Data Fig. 1:**
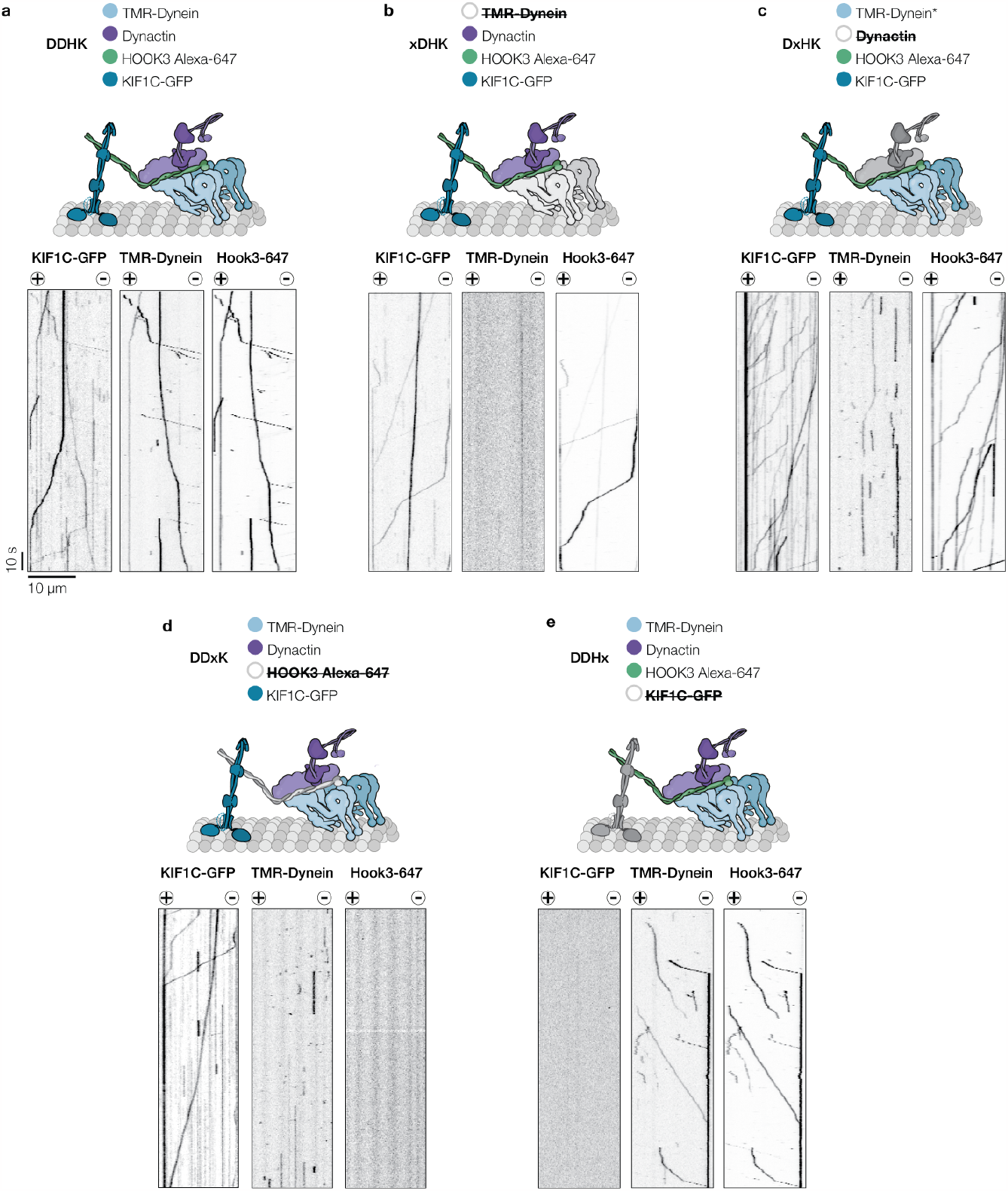
Controls for single molecule experiments. Representative kymographs from single molecule motility assays of Dynein-Dynactin-Hook3-KIF1C (DDHK) complexes showing (a) the full DDHK complex, and omission of individual components with (b) TMR-dynein omitted (xDHK), (c) dynactin (DxHK) and (d) HOOK3-Alexa 647 (DDxK) and (e) KIF1C-GFP (DDHx). The omitted factor is shown in grey in the cartoons and crossed out in the component list. Microtubule polarity is indicated with (+) and (-) signs. Note that motility of KIF1C and Dynein both towards the plus and minus end of microtubules is only observed in the complete DDHK complex.

**Extended Data Fig. 2:**
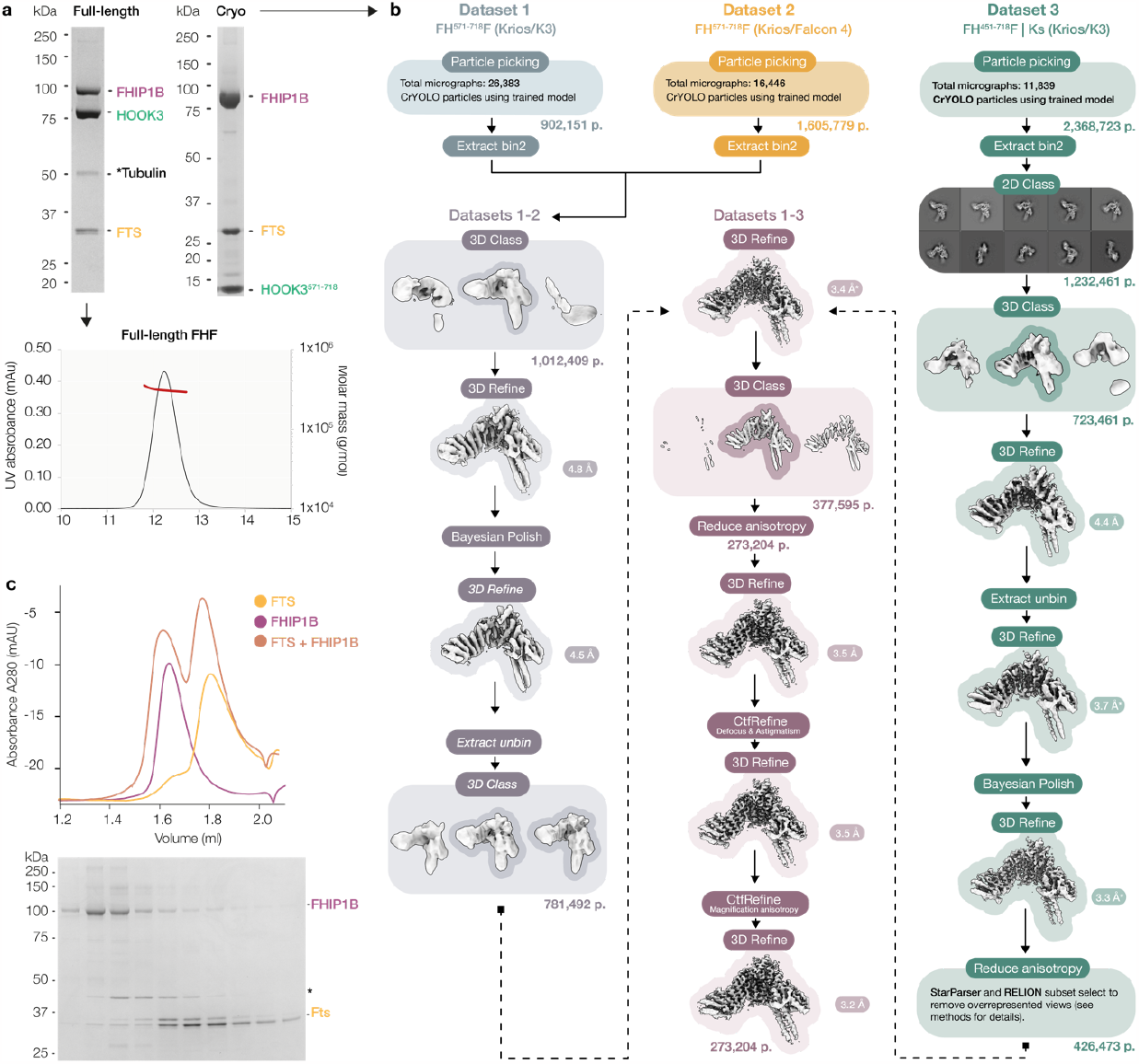
Biochemical analysis of FHF complex formation and FHF cryo-EM processing workflow. (a) SDS-PAGE of the full-length (left) FHF complex and corresponding SEC-MALS chromatogram (below gel), revealing an expected molecular weight of ~304 kDa. On the left is one of the truncated FHF constructs used for cryo-EM structure determination (containing HOOK3 residues 571-718). (b) Cryo-EM processing workflow for three combined FHF datasets. All steps were performed in RELION-4.1^64^ apart from particle picking, which was performed in CrYOLO^65^. Note that dataset 3 contained a construct of KIF1C stalk (GST-KIF1C stalk 674-922) but this did not alter the overall conformation of FHF as confirmed by comparison to FHF only datasets. Resolution values refer to post-sharpened and masked maps using the PostProcessing step of RELION. (c) Chromatogram of isolated 16 µM FTS, isolated 8 µM FHIP1B and FTS+FHIP1B added together. SDS-PAGE gel shows bands relevant to the combined FTS+FHIP1B run.

**Extended Data Fig. 3:**
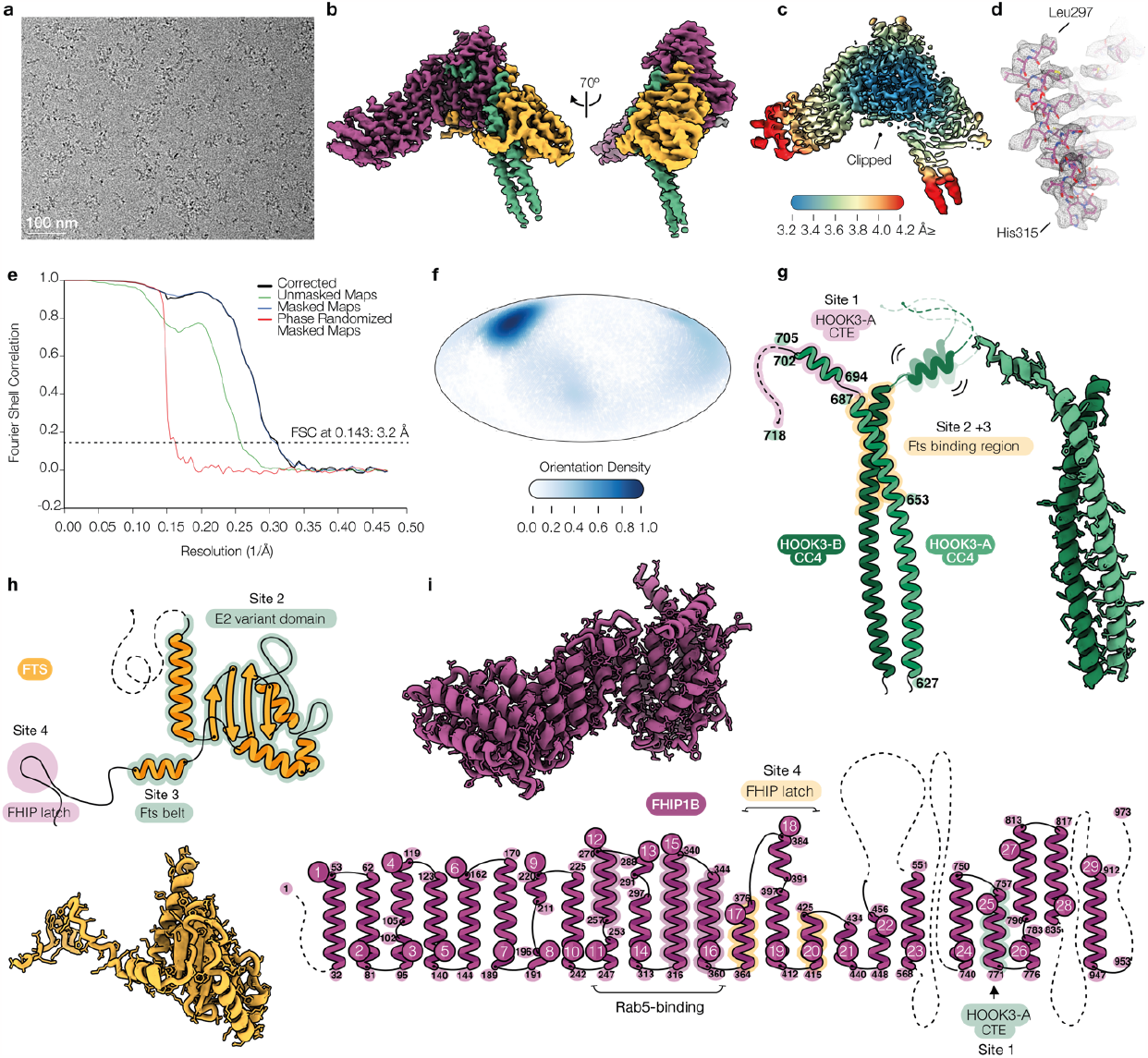
Validation and structural detail of FHF cryo-EM structure. (a), Integrated and motion corrected micrograph of FHF complexes from movie collected using a K3 camera in counting mode (1.09 Å/pixel). (b) Segmented cryo-EM density of FHF complex shown in front and side views. (c) RELION local-resolution plot applied to a clipped front view of the FHF complex. d, Example cryo-EM density (shown in mesh) from FHIP1B alpha helix (297-315) and corresponding built model. (e) Gold standard Fourier shell correlation (FSC) curves as determined by RELION-4.1 (FSC=0.143). (f) 2D representation of angular distribution extracted from particles.star of the final structure. (g-i) Topology and secondary structure of (g) HOOK3, (h) FTS and (i) FHIP1B. Interacting regions with other FHF components labelled and highlighted in transparent colour.

**Extended Data Fig. 4:**
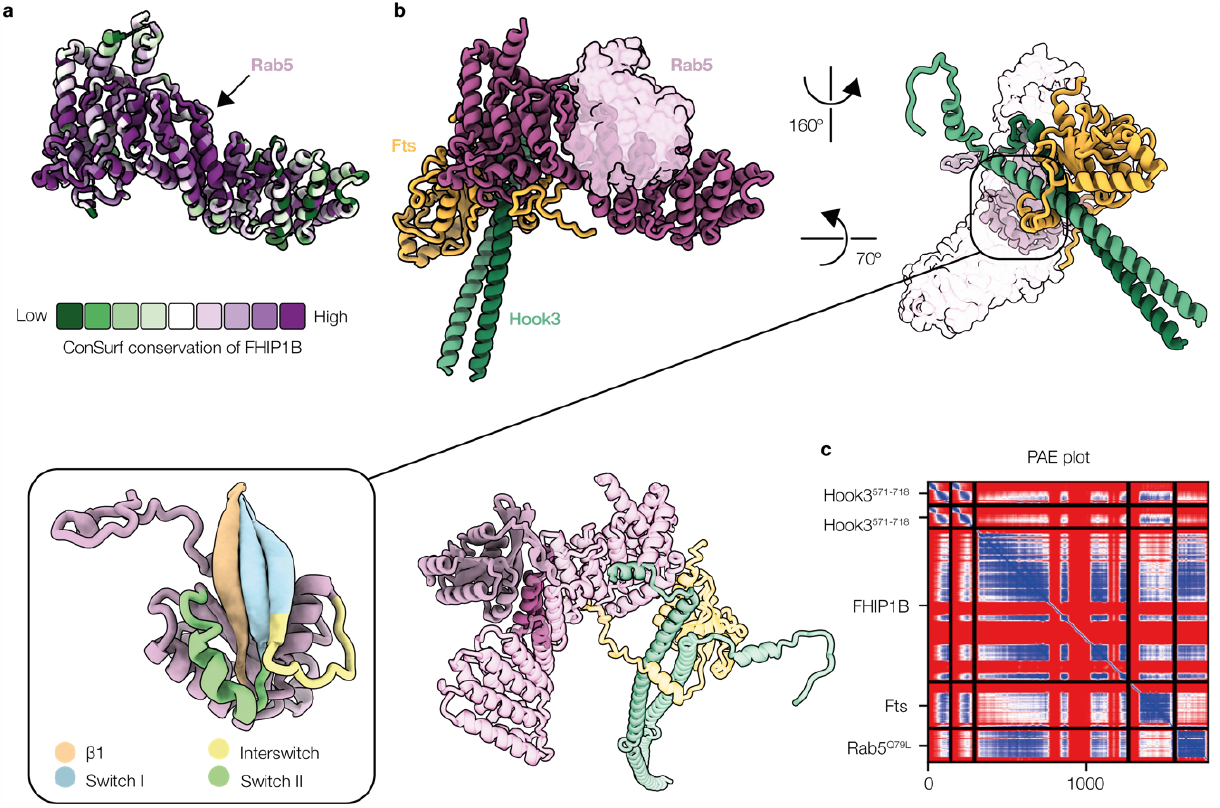
Conservation of FHIP1B and putative binding to Rab5. (a) ConSurf^78^ evolutionary conservation profile of FHIP1B. Model of FHIP1B coloured according to level of conservation (key below) and the view of the structure is the same as that in (b). (b) AlphaFold2 prediction of FH^571-718^F+Rab5^Q79L^. Predicted structure was superimposed on experimental cryo-EM FHF structure and used to show the interaction with Rab5 in three different views (top row middle and right panels, and bottom row, middle panel). Boxed inset shows predicted domain boundaries of Rab5 based on^79^. (c) AlphaFold2 Predicted Aligned Error (PAE) plot of FH^571-718^F+Rab5^Q79L^.

**Extended Data Fig. 5:**
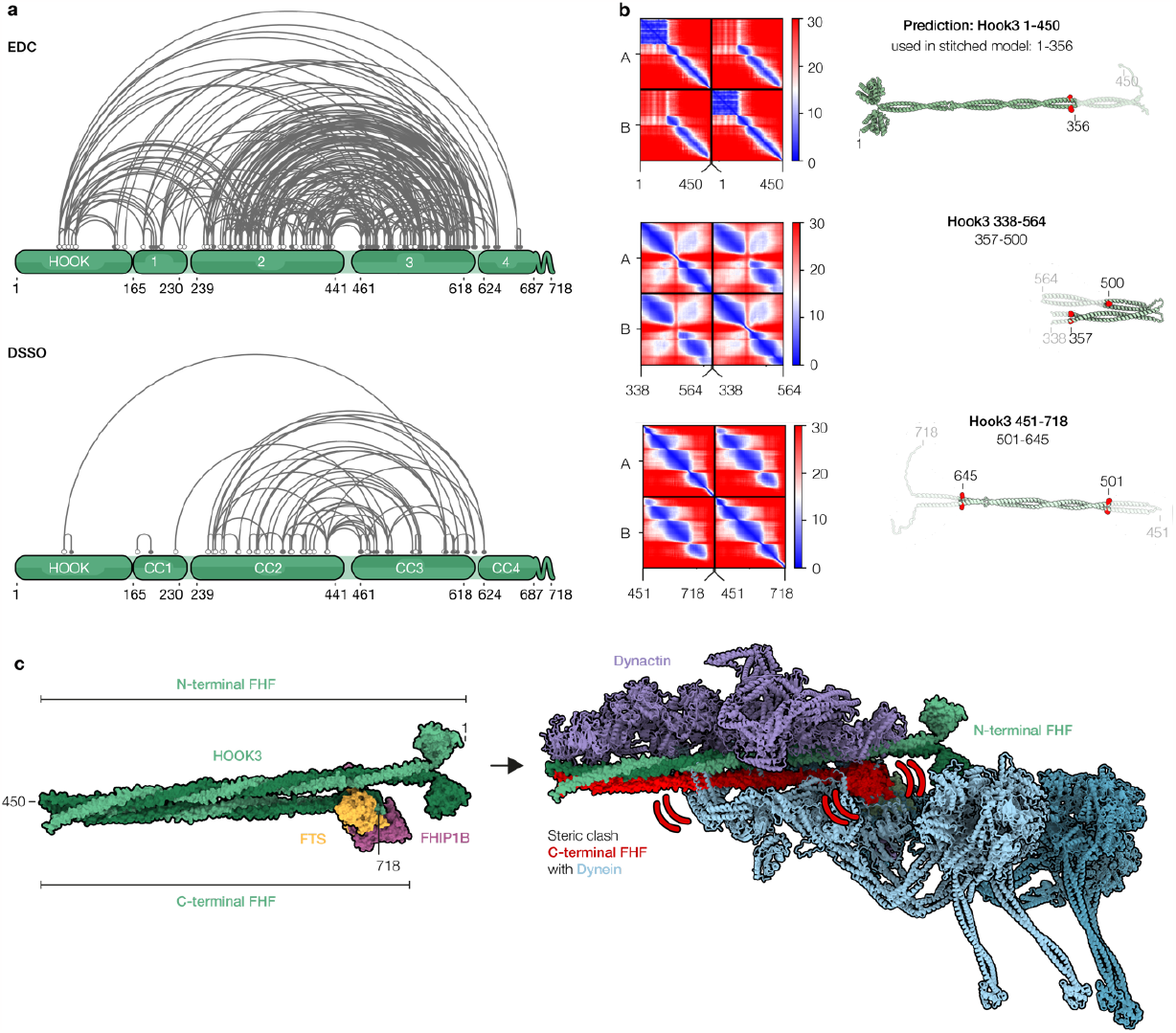
Crosslinking mass spectrometry and structural modelling of the auto-inhibited state of FHF. (a) Intramolecular crosslinks mapped on the full-length HOOK3 molecule, using EDC (top) and DSSO (bottom) crosslinkers. (b) AlphaFold2 Predicted Aligned Error (PAE) plots and corresponding structural prediction of three partially overlapping HOOK3 fragments (lengths of fragments indicated in the figure). Fragments were aligned at overlapping regions using the MatchMaker tool in UCSF Chimera^80^ regions of overlap were deleted at the regions indicated by the red spheres shown on the right. (c) The stitched full-length model of FHF (shown in surface rendering) predicts a steric clash upon binding to the dynein-dynactin complex (shown in cartoon rendering). In particular, the C-terminus of HOOK3, FTS and FHIP1B within auto-inhibited FHF is incompatible with stable binding to the dynein tails.

**Extended Data Fig. 6:**
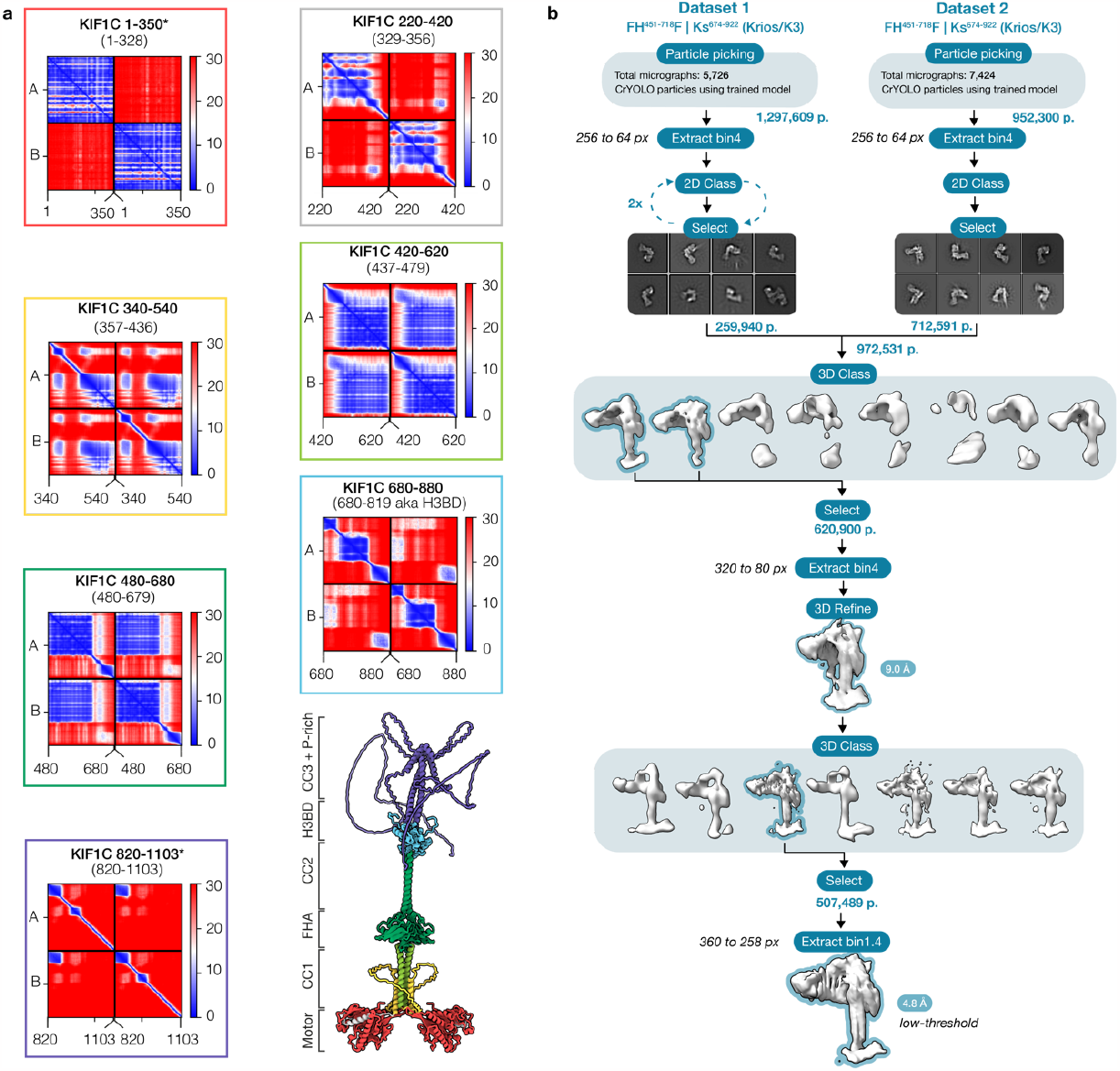
Structural modelling of full-length KIF1C and cryo-EM processing workflow for FH451-718F bound to KIF1C stalk (674-922). (a) AlphaFold2 Predicted Aligned Error (PAE) plots of KIF1C segments used to create the stitched model of the full-length molecule. Asterisk (*) in 1-350 prediction refers to use of only the monomer motor being used for superposition onto the 220-420 dimer structure. * in 820-1103 prediction highlights the high level of disorder predicted for the Proline-rich region (this segment is used for completion of structure only rather than for analysis). Sequences in brackets are the residues left after removal of overlapping superposed regions within the stitched molecule. (b) Cryo-EM processing workflow for two combined FHF-KIF1C stalk datasets. RELION-4.1^64^ was used throughout, apart from particle picking which was performed in crYOLO^65^.

**Extended Data Fig. 7:**
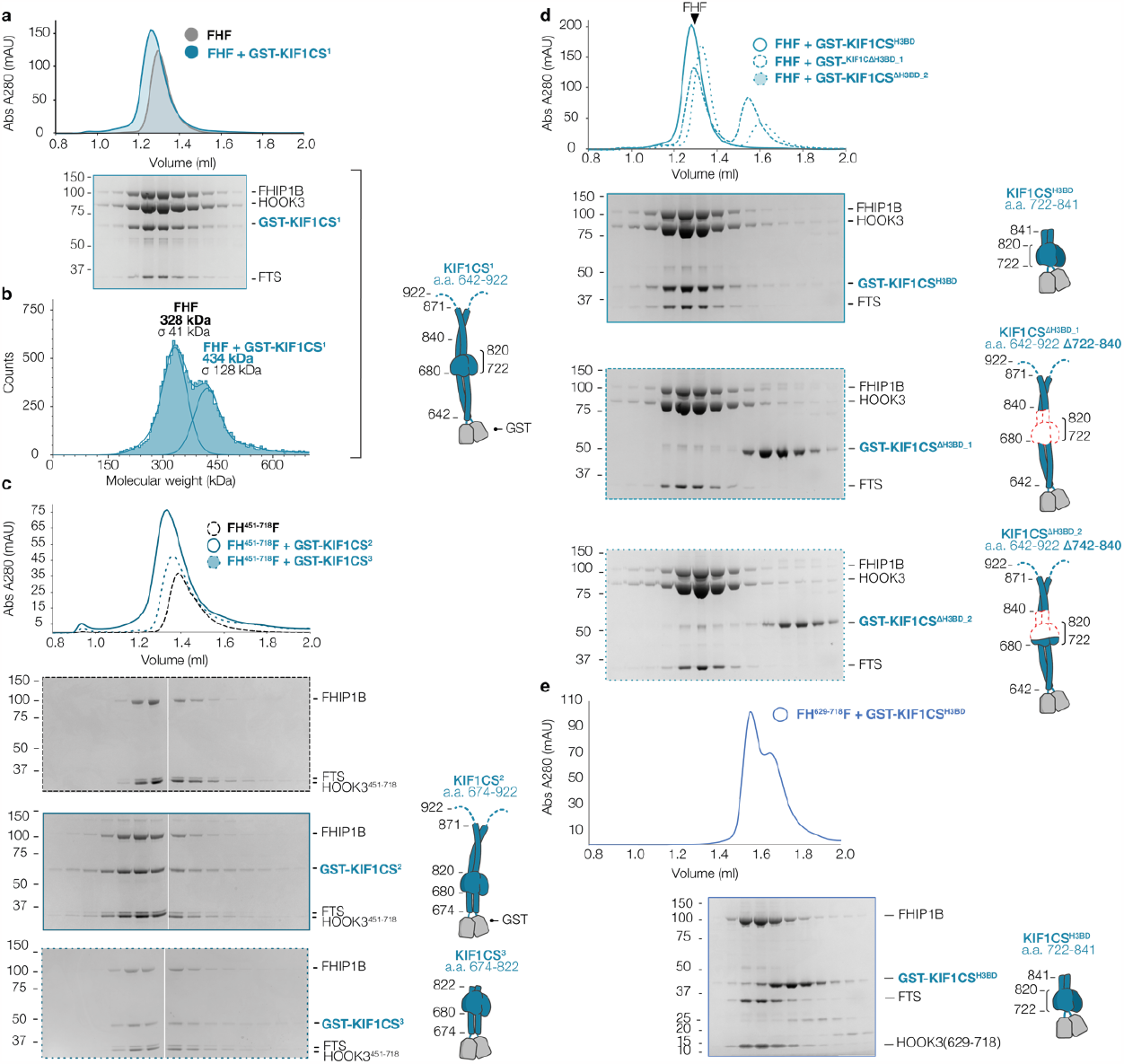
Biochemical characterisation of the FHF and KIF1C stalk interaction. (a) Size Exclusion Chromatography (SEC) of isolated full-length FHF and FHF+KIF1C stalk (residues 642-922). (b) Mass photometry of full-length FHF and KIF1C stalk using the Refeyn machine. Cartoon on the right depicts the KIF1C stalk construct used for a-b. (c) SEC reconstitution of FH^451-718^F in the absence or presence of two shorter KIF1C stalk constructs (residues 674-922 and 674-822, respectively) depicted in the cartoons on the right. (d) SEC reconstitution of full-length FHF in the presence and absence of the KIF1C stalk H3BD domain. Two H3BD deletions constructs within KIF1C stalk were tested (∆722-840 and ∆742-840) against H3BD construct (residues 722-841), as depicted in the cartoons on the right. (e) SEC reconstitution of truncated FH^629-718^F with H3BD shows no binding. SDS-PAGE gels for all runs show fractions from the peak regions.

**Extended Data Fig. 8:**
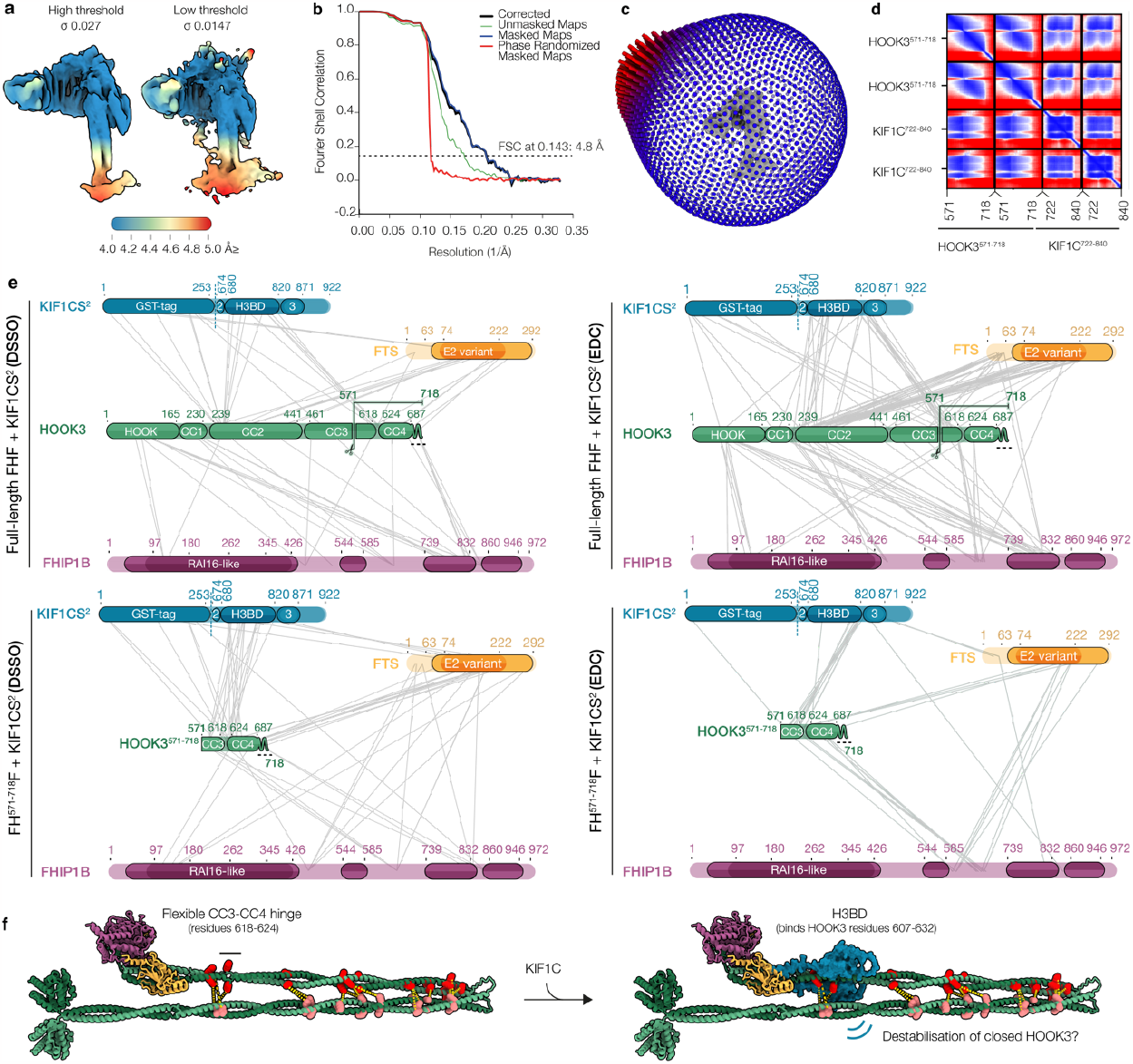
FHF-KIF1C cryo-EM, AlphaFold2 and cross-linking mass spectrometry validation and analysis. (a) RELION local-resolution plot applied to the FH^451-718^F-KIF1CS^674-922^ cryo-EM structure, shown at two different density thresholds. (b) Gold standard Fourier shell correlation (FSC) curves as determined by RELION-4.1 (FSC=0.143). (c) 3D depiction of angular distribution for the final FH^451-718^F-KIF1CS^674-922^ structure with the cryo-EM density shown in the middle. (d) AlphaFold2 Predicted Aligned Error (PAE) plot for the HOOK3^571-718^-H3BD^722-840^ prediction. (e) Crosslinking mass spectrometry results for full-length FHF or FH^571-718^F bound to KIF1C stalk (containing residues 674-922), either using the EDC or DSSO crosslinkers. (f) Intramolecular interactions within the predicted model of auto-inhibited FHF could be partially disrupted by the KIF1C H3BD domain binding at the CC3-CC4 hinge at close proximity to the HOOK3 CC3-CC4 intramolecular interaction with CC2.

**Extended Data Fig. 9:**
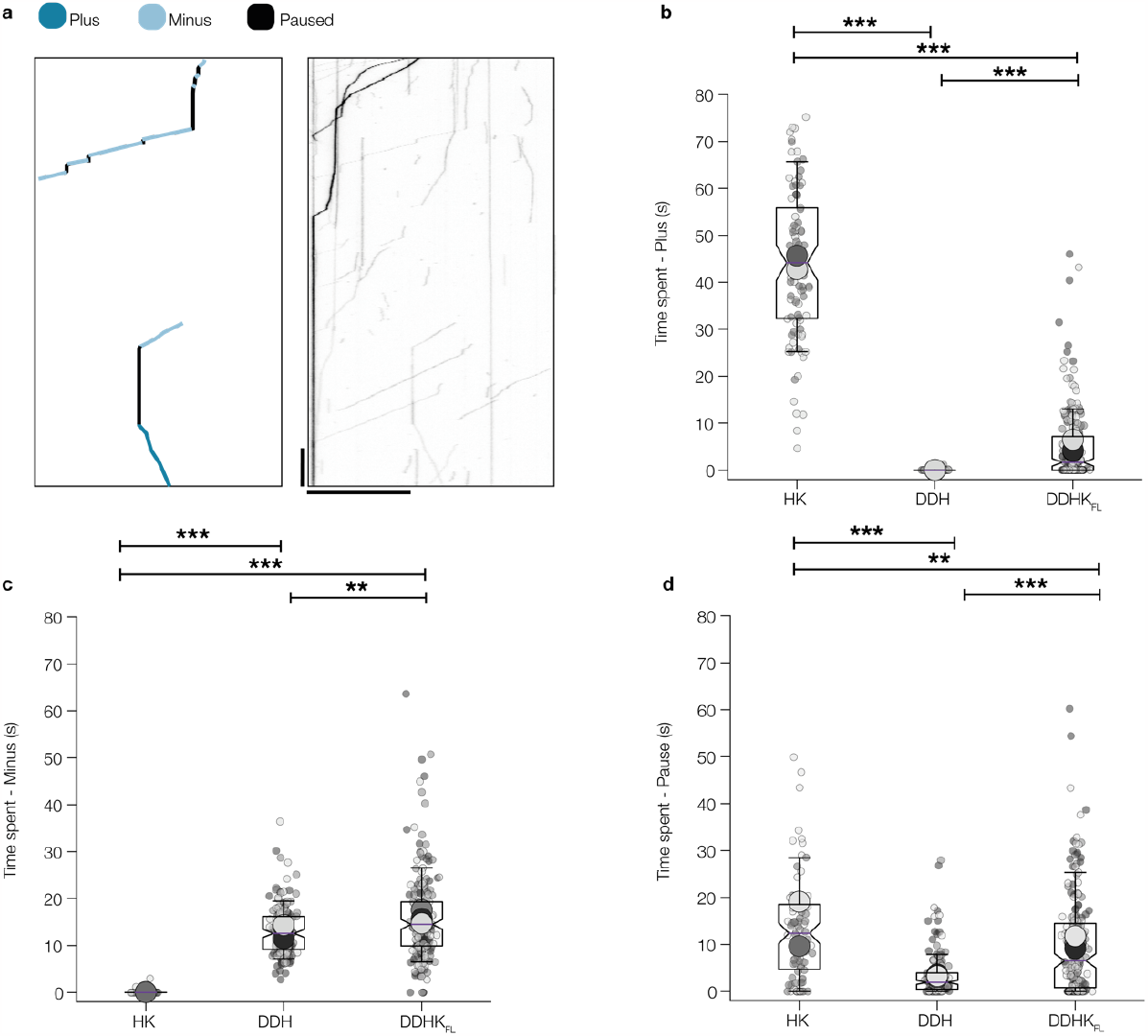
Subclassification of motor complex motility and pausing events. (a) Example kymograph and annotation showing what would be classified as plus and minus end directed motility and paused (moving < 25nm/s). The example shows the TMR-dynein channel of a DDHK chamber. The scale bar is 20 μm and 20 s in the horizontal and vertical axes, respectively. (**b-d**) Superplots of HK, DDH and DDHK time spent moving towards the plus and minus ends of the microtubule, or spent pausing at a speed < 25nm/s. Note that wholly static tracks (total distance <1000 nm) are excluded from this analysis. Small dots show data per microtubule and large dots show experimental averages. Boxes show quartiles with whiskers spanning 10%-90% of the data. n=94-202 microtubules. **p<0.01, ***p<0.001 using Kruskal Wallis H test followed by a Conover’s posthoc test to evaluate pairwise interactions with a multiple comparison correction applied using Holm–Bonferroni.

**Extended Data Fig. 10:**
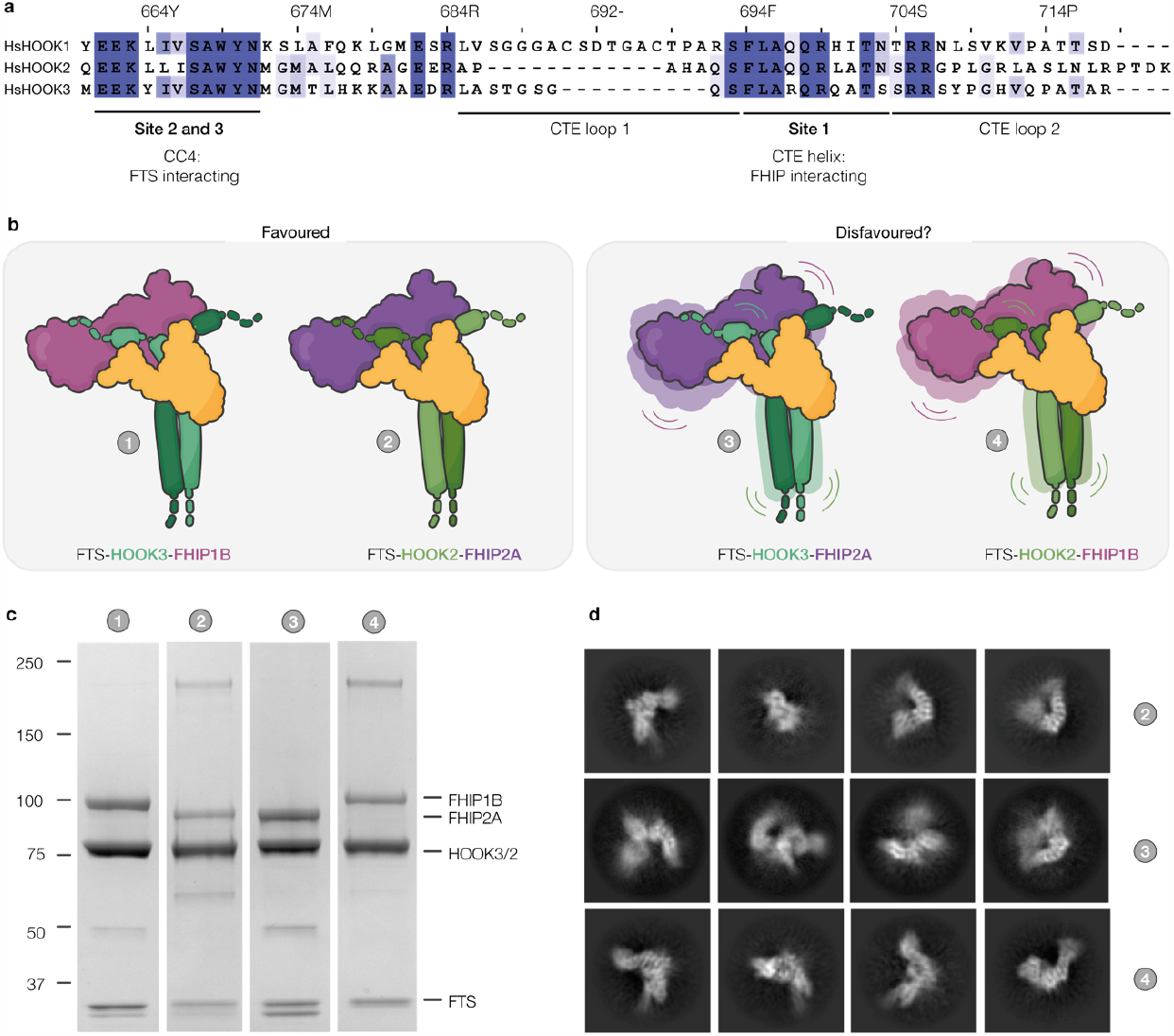
HOOK specificity for different FHIPs within FHF is unlikely to be dictated by sequence or structure. (**a**) Sequence conservation between human HOOK1, 2 and 3. Residue numbers refer to HOOK3. (**b**) The combinations of FHF co-expressed and purified to test if complex formation forms. FTS-HOOK3-FHIP1B (complex 1) and FTS-HOOK2-FHIP2A (complex 2) classed as favoured assemblies and FTS-HOOK3-FHIP2A (complex 3) and FTS-HOOK2-FHIP1B (complex 4) are classed as disfavoured assemblies based on^34^. (**c**) SDS-PAGE purified FHF (complexes 1-4) show that HOOK2 or HOOK3 promiscuously bind FHIP2A or FHIP1B in vitro. (**d**) Cryo-EM 2D class averages of favoured (2) and disfavoured (3 and 4) complexes show recognisable FHF views in all cases, indicating stable FTS-HOOK-FHIP assembly in all combinations tested. Extended Data

**Extended Data Table 1:**
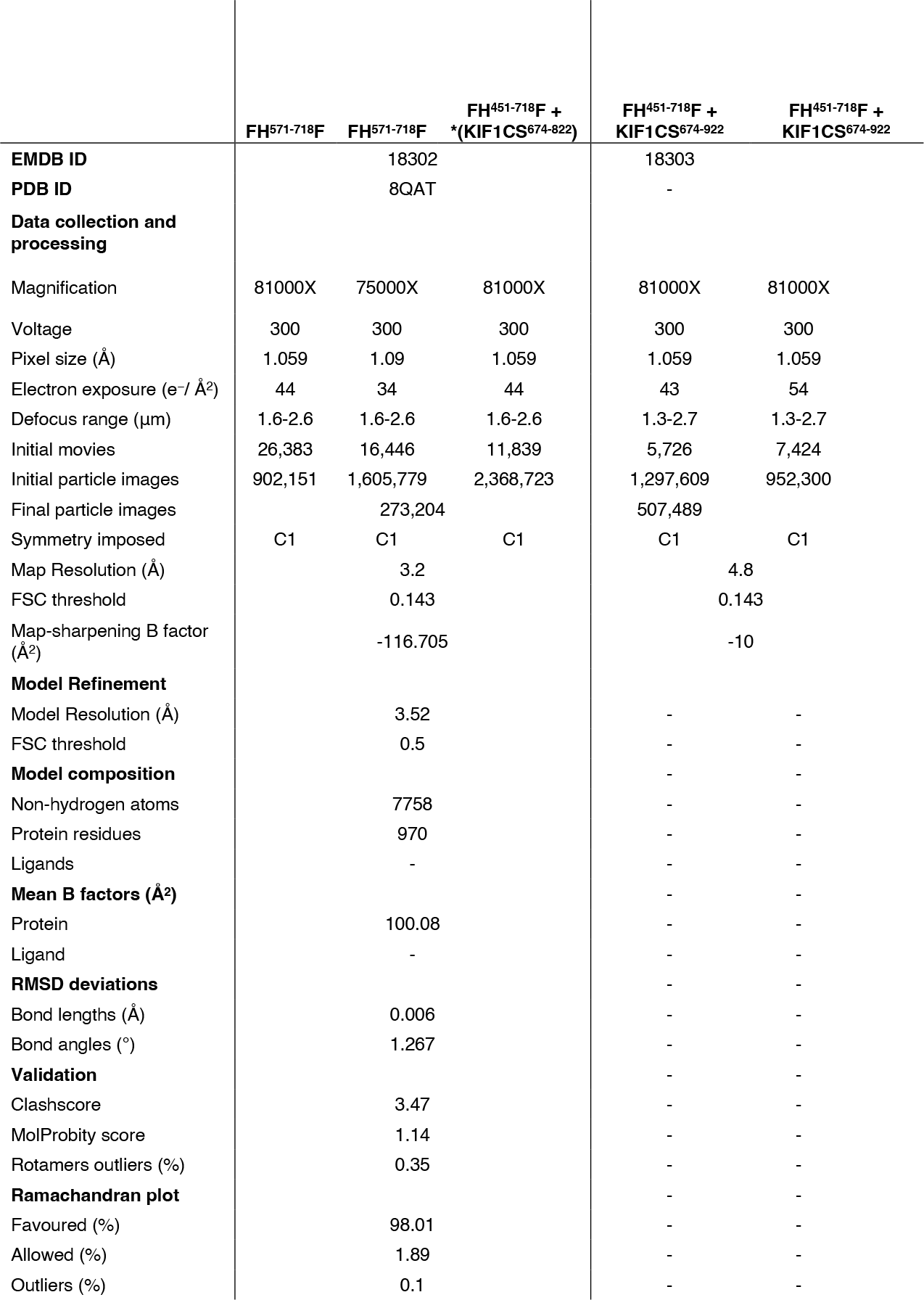
Cryo-EM and refinement statistics.

